# Mechanostat parameters estimated from time-lapsed *in vivo* micro-computed tomography data of mechanically driven bone adaptation are logarithmically dependent on loading frequency

**DOI:** 10.1101/2023.01.07.523082

**Authors:** Francisco C. Marques, Daniele Boaretti, Matthias Walle, Ariane C. Scheuren, Friederike A. Schulte, Ralph Müller

## Abstract

Mechanical loading is a key factor governing bone adaptation. Both preclinical and clinical studies have demonstrated its effects on bone tissue, which were also notably predicted in the mechanostat theory. Indeed, existing methods to quantify bone mechanoregulation have successfully associated the frequency of (re)modeling events with local mechanical signals, combining time-lapsed *in vivo* micro-computed tomography (micro-CT) imaging and micro-finite element (micro-FE) analysis. However, a correlation between the local surface velocity of (re)modeling events and mechanical signals has not been shown. As many degenerative bone diseases have also been linked to impaired bone (re)modeling, this relationship could provide an advantage in detecting the effects of such conditions and advance our understanding of the underlying mechanisms. Therefore, in this study, we introduce a novel method to estimate (re)modeling velocity curves from time-lapsed *in vivo* mouse caudal vertebrae data under static and cyclic mechanical loading. These curves can be fitted with piecewise linear functions as proposed in the mechanostat theory. Accordingly, new (re)modeling parameters can be derived from such data, including formation saturation levels, resorption velocity modulus, and (re)modeling thresholds. Our results revealed that the norm of the gradient of strain energy density yielded the highest accuracy in quantifying mechanoregulation data using micro-FE analysis with homogeneous material properties, while effective strain was the best predictor for micro-FE analysis with heterogeneous material properties. Furthermore, (re)modeling velocity curves could be accurately described with piecewise linear and hyperbola functions (root mean square error below 0.2 µm/day for weekly analysis), and several (re)modeling parameters determined from these curves followed a logarithmic relationship with loading frequency. Crucially, (re)modeling velocity curves and derived parameters could detect differences in mechanically driven bone adaptation, which complemented previous results showing a logarithmic relationship between loading frequency and net change in bone volume fraction over four weeks. Together, we expect this data to support the calibration of *in silico* models of bone adaptation and the characterization of the effects of mechanical loading and pharmaceutical treatment interventions *in vivo*.

## Introduction

Bone is a dynamic organ capable of adapting to its mechanical environment (Burr et al. 2002). Through a multiscale process, loads are transferred from the organ to the cellular level, eliciting highly coordinated biological responses that adapt its architecture (Wolff 1986, Lanyon 1992, Turner 1998). This process of continuous bone formation and resorption comprises both bone modeling and remodeling events, collectively referred to as (re)modeling (Huiskes et al. 2000). Indeed, several studies have successfully shown the influence of mechanical loading in bone adaptation, especially in trabecular bone, highlighting how mechanical cues guide the bone structure towards an optimal load transfer (Lanyon 1992, Rubin and McLeod 1994, Huiskes et al. 2000, De Souza et al. 2005, Lambers et al. 2011). From a mathematical modeling perspective, the mechanostat theory (Frost 1987) is a widely known proposal for the regulatory mechanism driving tissue-level bone adaptation. Mirroring the function of a thermostat, a system to maintain the temperature within a predefined interval through heating and cooling, Frost argued that bone physiology also embodies processes to maintain bone mass within an adapted window for optimal mechanical usage. Specifically, Frost postulated the existence of key set-points or thresholds that bound this adapted window such that bone would modulate states of disuse-remodeling, i.e., when mechanical demands are lower than normal named “disuse”, or bone modeling, i.e., when mechanical demands are higher than normal termed “mild overload”, where bone mass is predominantly added or removed, accordingly. This fine balance would eventually normalize the mechanical demands perceived on the structure to the adapted window. Additionally, Frost reflected on the mechanostat theory in the context of disease states, such as osteoporosis and osteogenesis imperfecta, space orbit, or in the presence of circulating agents, like growth factors or medications, and proposed how these would interfere with the physiological mechanoregulation of the adapted window. Briefly, he associated some disease states with a desensitization of response mechanisms, which would raise bone modeling thresholds, hence reducing the adaptive response when the mechanical usage increased, while high mechanically demanding conditions would lower such thresholds, making the system overreactive and triggering strong anabolic reactions. Crucially, the mechanostat theory has been successfully incorporated into *in silico* models of bone adaptation capable of approximating the trends observed *in vivo* in response to various interventions (Levchuk et al. 2014) and loading frequencies (Kameo et al. 2011). Such models can leverage time-lapsed *in vivo* micro-computed tomography (micro-CT) data, which has also enabled tracking structural changes in response to externally applied loading in preclinical animal studies, contributing to a comprehensive description of both morphometric changes and mechanoregulation information of bone adaptation (Schulte et al. 2013, Birkhold et al. 2017, Albiol et al. 2020, San Cheong et al. 2020).

Notably, with aging and in specific disease contexts, this (re)modeling process becomes unbalanced (Rubin et al. 1992, Bassey et al. 1998), often leading to other degenerative conditions such as osteoporosis, which precede an increased risk of fractures and culminate in considerable health and economic costs for society (Gabriel et al. 2002, Becker et al. 2010). In agreement with Frost’s assumptions, impaired bone mechanoregulation has been suggested as a possible cause of this problem. Therefore, advancing our ability to retrieve mechanoregulation information from time-lapsed *in vivo* micro-CT data can help better understand the underlying mechanisms and, with that, develop more effective treatments for these degenerative conditions.

In this regard, preclinical models, such as the mouse caudal vertebrae or tibia loading model, have been foundational to explore bone adaptation processes by enabling controllable experimental settings that can also mimic clinical pathological conditions (Rubin and McLeod 1994, Robling and Turner 2002, Webster et al. 2008, Vandamme 2014, Razi et al. 2015, Roberts et al. 2019). As a result, existing methods to investigate mechanoregulation have successfully linked (re)modeling events with tissue strains obtained from micro-finite element (micro-FE) analysis, showing strong associations between formation/resorption and high/low tissue strains, respectively (Schulte et al. 2013, Razi et al. 2015, Scheuren et al. 2020), which were summarized in conditional probability curves defined over a continuous range of tissue strains. To this end, several mathematical quantities have been proposed to describe the local mechanical signal: strain energy density (SED), effective strain (Pistoia et al. 2002), and norm of the gradient of SED (∇SED). Deformation can be quantified using SED or effective strain, a derived quantity from SED that accounts for differences in tissue Young’s modulus. Likewise, it is hypothesized that load-induced bone adaptation arises from mechanical deformation perceived by osteocytes, the mechanosensitive cells in bone (direct cell strain), and interstitial fluid flow (shear stress) in the lacunar-canalicular network (Klein-Nulend et al. 1995, Fritton and Weinbaum 2009, Weinbaum et al. 2011). Mathematically, ∇SED represents spatial differences in tissue deformation, which are believed to induce fluid flow (Huiskes 2000). From this perspective, it is unclear which mechanical signal shows the best association with bone (re)modeling events.

Furthermore, conditional probability-based approaches lack quantitative information describing the expected change in bone material for a given strain value. This dependency has been considered *in silico* (Adachi et al. 2001, Levchuk et al. 2014, Goda et al. 2016, Louna et al. 2019) by defining the normal surface velocity in response to the perceived mechanical signals guiding (re)modeling events. This formulation based on surface velocity deviates from the original mechanostat theory by Frost, which focused on changes in bone mass with the mechanical signal. Nonetheless, this shift was vital for the development of such high-resolution *in silico* models based on micro-CT images driven by surface advection (Adachi et al. 2001, Levchuk et al. 2014), where new bone material is added along the gradient of the surface.

Besides, previous work has shown a dose-dependent effect of this mechanical stimulus (Mosley and Lanyon 1998, Sugiyama et al. 2012, Scheuren et al. 2020, Walle et al. 2021), but an association of surface velocity with mechanical signal has not been investigated at the tissue level *in vivo*. On the one hand, it hinders more detailed comparisons of degenerative conditions that may conserve the mechanosensation ability of bone cells, associated with mechanical signal thresholds that regulate (re)modeling events but influence the magnitude of the response to the mechanical stimuli and the overall bone turnover. On the other hand, from a computational modeling perspective, retrieving such information from *in vivo* data can provide valuable calibration data for the mechanoregulation mechanisms implemented in such models to achieve more realistic bone adaptation representations.

Therefore, the present study had two aims. First, we sought to identify which mechanical signal showed the best association with bone (re)modeling events using conditional probability curves and quantified with the correct classification rate (CCR) (Tourolle né Betts et al. 2020). Second, we aimed to propose a method to retrieve mechanoregulation information from time-lapsed *in vivo* micro-CT data that associates the surface (re)modeling velocity (RmV) with the local mechanical signal. We express this relationship in RmV curves from which several biologically meaningful parameters can be derived, such as formation/resorption thresholds and saturation levels. Notably, a consistent nomenclature of these parameters is proposed and formalized in alignment with the current understanding of the mechanostat theory. Furthermore, we applied our novel method to an *in vivo* mouse caudal vertebrae dataset (Scheuren et al. 2020) and quantified the local dynamic response of trabecular bone adaptation to static and cyclic loading of varying frequencies. We hypothesized that the effect of increased loading frequencies could be measured with our new analysis and compared through the parameters derived from these RmV curves. Finally, we investigated if there was a relationship between these parameters and loading frequency, analogous to the logarithmic relationship observed between loading frequency and net change in bone volume fraction over the 4-week observation period (Scheuren et al. 2020).

## Materials and Methods

### Time-lapsed *in vivo* micro-CT mouse caudal vertebrae dataset

The experimental data used for this study was collected in a previous longitudinal murine *in vivo* loading study (Scheuren et al. 2020), supporting 3R principles. Briefly, 11-week-old female C57BL/6J mice received surgery to allow mechanical loading of the sixth caudal vertebrae (CV6) via stainless steel pins (Fine Science Tools, Heidelberg, Germany) inserted into the fifth and seventh vertebrae following the protocol by Webster et al. (2008). After surgery and recovery, the 15-week-old mice were split into five groups: sham loading (0 N, 8 mice), 8 N static (8 mice), or 8 N cyclic loading with the frequencies of 2 Hz (7 mice), 5 Hz (5 mice), or 10 Hz (8 mice). The loading regime was performed for five minutes, three times per week, over four weeks, as previously described by Lambers et al. (2011). Cyclic loading aimed at recreating supraphysiological loading conditions, that is, which surpass the loading perceived with typical physiological activity of the mice (Frost 2003). With the start of loading, the animals were scanned weekly using *in vivo* micro-CT (vivaCT 40, Scanco Medical AG, Switzerland), with an isotropic voxel size of 10.5 μm. All mouse experiments described in the present study were carried out in strict accordance with the recommendations and regulations in the Welfare Ordinance (TSchV 455.1) of the Swiss Federal Food Safety and Veterinary Office (license number 262/2016).

### Automated compartmental analysis of the mouse caudal vertebrae

Consecutive time-points of the micro-CT scans were initially registered to each other using the Image Processing Language (IPL Version 5.04c; Scanco Medical AG, Switzerland). For the identification of the trabecular and cortical compartments in the structure, the images were filtered with a Gaussian filter (sigma: 1.2, truncate: 1) as implemented in (Virtanen et al. 2020), binarized with a threshold of 580 mgHA/cm^3^ (Scheuren et al. 2020), followed by automatic identification of the relevant compartments following the protocol proposed by Lambers et al. (2011). This approach was implemented in Python (version 3.9.9) and validated against the existing pipeline in IPL (Supplementary Material 1.1.1 and Supplementary Figure 1).

### Micro-finite element analysis

Micro-CT images were analyzed with micro-finite element analysis (micro-FE) to estimate the local mechanical signal. Image voxels were converted to 8-node hexahedral elements (approximately 20 million elements in total), and bone was assumed to behave within the linear elastic regime. The nodes at the proximal end of the micro-FE mesh were constrained in all directions, while the nodes at the distal end were displaced by 1% of the length in the z-axis (longitudinal axis of the sample). Two sets of simulations were performed for each sample: with homogeneous and heterogeneous material properties based on the binary and grayscale images of the samples. The former considered a Young’s modulus value of 14.8 GPa for bone and 2 MPa for marrow (defined as non-bone voxels in the trabecular region) and a Poisson’s ratio of 0.3 (Webster et al. 2008) (see Supplementary Figure 2 for a sensitivity analysis of the effect of the Young’s modulus value assigned to bone and marrow voxels on the ability to quantify mechanoregulation information). Conversely, the latter applied a linear relationship between bone mineral density and Young’s modulus (Mulder et al. 2007) for all voxels within the outer mask of the vertebrae (closed mask that contains all voxels within the external surface of the vertebrae), also using a Poisson’s ratio of 0.3. In this case, the minimum value allowed for Young’s modulus was also 2 MPa. Two cylindrical discs were added at the proximal and distal ends of the vertebra model (Webster et al. 2008), mimicking the role of the intervertebral discs. Disc settings were calibrated for micro-FE with homogeneous and heterogeneous material properties (Supplementary Material 1.2). The simulations computed strain energy density (SED, in MPa) in the vertebrae, from which all derived quantities were determined after linear rescaling to match the forces applied in vivo: 8 N for loaded groups and 4 N (physiological loading) for the sham-loaded group (0 N) (Christen et al. 2012). The pipeline was also implemented in Python, and the simulations ran on the Euler cluster operated by Scientific IT Services at ETH Zurich, using the micro-FE solver ParOSol (Flaig and Arbenz 2011) on Intel Xeon Gold 6150 processors (2.7-3.7 GHz).

### Mechanoregulation analysis based on conditional probability curves

The mechanoregulation analysis performed in this study considered three mathematical quantities representing the local mechanical signal: strain energy density (SED), effective strain (Pistoia et al. 2002), and the norm of the gradient of SED (∇SED). In this context, the gradient was computed using the central difference scheme, and the norm was used as a proxy for the fluid flow in each voxel. For SED and effective strain, the values were collected on the voxels on the bone side of the surface interface between bone and marrow, while the values for ∇SED were collected on the marrow side. The latter was motivated by previous results showing that fluid flow velocity surrounding osteocytes was most influential close to the bone surfaces (Kameo et al. 2008) and, similarly, that fluid flow affected the activity of osteoblasts and mesenchymal stem cells (Riehl et al. 2015, Riehl et al. 2017), which are also primarily found on the bone surfaces. The analysis only considered the surface voxels and mechanical signal values inside the trabecular compartment identified with the algorithm described in a previous section.

The conditional probabilities of a (re)modeling event (formation, quiescence, resorption) to occur at each value of mechanical signal were calculated as described previously by Schulte et al. (2013), for weekly intervals (e.g., weeks 1-2 considered the micro-CT images from weeks 1 and 2 and the micro-FE from week 1) and the 4-week observation period (using the micro-CT images from weeks 0 and 4 and micro-FE data from week 0). Briefly, for each (re)modeling event, the values of the mechanical signal at the surface voxels were collected and normalized by the 99^th^ percentile of all observed values, defining a normalized mechanical signal (%). These values were binned at 1% of the 99^th^ percentile determined, and a normalization of the total number of counts per (re)modeling event was applied to rule out any dependence on the imbalance between these events before calculating conditional probabilities. A conditional probability of 0.33 indicates an equal probability of any (re)modeling event occurring.

The quantification of the amount of mechanoregulation information recovered in the analysis relied on the CCR, using an implementation proposed by Tourolle né Betts et al. (2020). This metric summarizes in a single number the accuracy of this ternary classification problem that categorizes (re)modeling events (formation, quiescence, and resorption) within the range of observed local mechanical signal values. In short, two thresholds (t_r_ and t_f_) define three intervals, one per (re)modeling event: resorption (mechanical signal < t_r_), quiescence (t_r_ < mechanical signal < t_f_), and formation (t_f_ < mechanical signal). A 3-by-3 confusion matrix is filled with the sum of the conditional probabilities of each event within the intervals: diagonal entries (e.g., the sum of the conditional probability values of formation within the formation interval) represent true positive entries, while off-diagonal are false negatives and positives. The normalized trace of the confusion matrix yields the CCR value (optimal thresholds are found by sweeping the range of mechanical signal values until the highest CCR is obtained). A value of 1 represents perfect accuracy, and 0.33 describes the accuracy of a random classification, given the presence of three classes. In this context, CCR values above 0.33 indicate that the mechanical signal descriptor considered can predict (re)modeling events with better accuracy than a random classifier.

### Mechanostat (re)modeling velocity curve and parameter derivation

Here, we introduce a novel method to estimate sample-specific (re)modeling velocity curves and their corresponding mechanostat parameters based on time-lapsed micro-CT data. The proposed method considers the scans from two time-points: baseline and follow-up images. Note that only the trabecular compartment is considered for this analysis. First, the follow-up image is registered to the baseline, revealing volumes of formed, quiescent, and resorbed clusters. Next, these clusters are used to classify surface voxels of the baseline image: formation surfaces consist of the overlap between dilated formed clusters and the baseline surface (von Neumann neighborhood), resorption surfaces refer to the overlap between resorbed clusters and the baseline surface, and quiescent surfaces contain the remaining surface voxels. A distance transform (DT) algorithm (taxicab metric) is applied to the follow-up image and the inverse of the follow-up image and masked with the formation and resorption surfaces identified before, yielding the distance of each surface voxel to the surface of the follow-up scan. It is assumed that the distance transform of the follow-up reveals the amount of formed bone, while the inverted follow-up identifies the depth of resorption per surface voxel. The values assigned to formation surfaces are obtained by gray-dilating (von Neumann neighborhood) the distance transform values of the formed clusters into the neighboring formation surface voxels identified. A gray-dilation operation is needed since the remodeling distance values are no longer binary (in comparison to the identification of remodeling surfaces performed initially) and to ensure an accurate distance calculation (e.g., if a 1-voxel thick is added on the surface, the surface voxels underneath will contain a remodeling distance value of 2, despite a single layer of voxels being added; with the gray dilation, the value of distance one computed by the distance transform on the added layer can be accurately projected to the surface voxels). Further, these are linearly scaled in a cluster-specific fashion to match the volume of the corresponding cluster (Supplementary Figure 3). Next, the mechanical signal (ms) computed from the micro-FE analysis of the baseline image is collected using the same (re)modeling surface masks. For this application, the micro-FE analysis considered the homogeneous material properties described previously. Furthermore, we selected effective strain in microstrain (µε) as the mechanical signal descriptor, although an alternative signal would also be suitable (e.g., SED or ∇SED).

Given that each surface voxel contains information on the amount of surface change and the estimated mechanical signal, a 2D histogram is computed, considering the mechanical signal on the horizontal axis and the estimated distance on the vertical axis. The mechanical signal is capped at the 99^th^ percentile to eliminate very high (unphysiological) values and binned at 1% of this value. In the vertical axis, all values are considered and binned at 1% of the maximum value observed. A weighted average of the distance values is computed using the number of counts in the 2D histogram as weights, providing a value for each mechanical signal bin and considering a value of 0 for the quiescent surface voxels. The last step converts the estimated distance to a (re)modeling velocity magnitude by multiplying and dividing by the voxel size of the images and the interval between the time-points analyzed, respectively. For consistency with other dynamic morphometry quantities (such as mineral apposition and resorption rates), we chose to express (re)modeling velocity in µm/day. In this context, (re)modeling velocity aims to describe the surface movement along the normal surface direction (Adachi et al. 2001, Levchuk et al. 2014, Goda et al. 2016, Louna et al. 2019).

Finally, mathematical functions are fitted to the curves obtained, yielding their quantitative parametric description, namely: piecewise linear (Equation 1), as proposed in the original mechanostat theory, and a continuous hyperbola function (Equation 2), both illustrated in Figure 1, and which enable quantifying new (re)modeling parameters *in vivo*. The piecewise linear function is defined by formation and resorption saturation levels (FSL and RSL, µm/day), which determine the maximum and minimum (re)modeling velocities observed, formation and resorption thresholds (FT and RT, µε) which determine the minimum and maximum mechanical signal value from which formation and resorption events are observed, and formation and resorption velocity modulus (FVM and RVM, µm/day / µε) which determine the change in (re)modeling velocity resulting from a change in mechanical signal, defined between FSL, RSL and FT, RT, respectively. Specifically, we highlight the proposal to define FVM and RVM as a modulus because the values are also proportional to mechanical strain. Comparably, the hyperbola function comprises similar FSL and RSL parameters, a (re)modeling threshold (RmT, µm/day) corresponding to the mechanical signal value at which the RmV curve is zero and a (re)modeling velocity modulus (RmVM, µm/day x µε), which is defined as the scale factor determining the rate of change in (re)modeling velocity resulting from a change in mechanical signal. A summary of these parameters is also presented in Table 1.

**Table 1.**
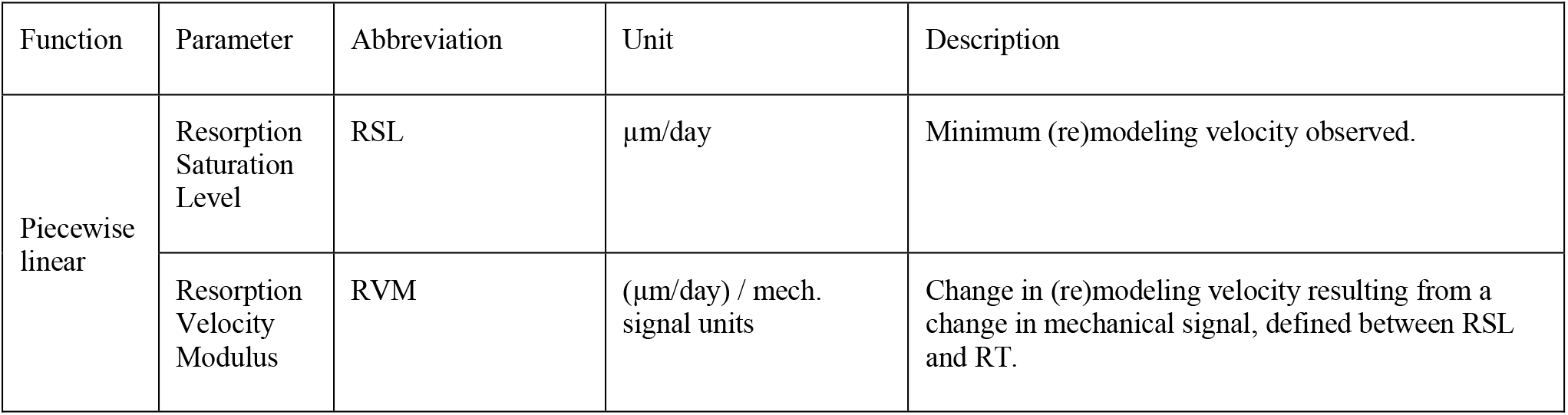
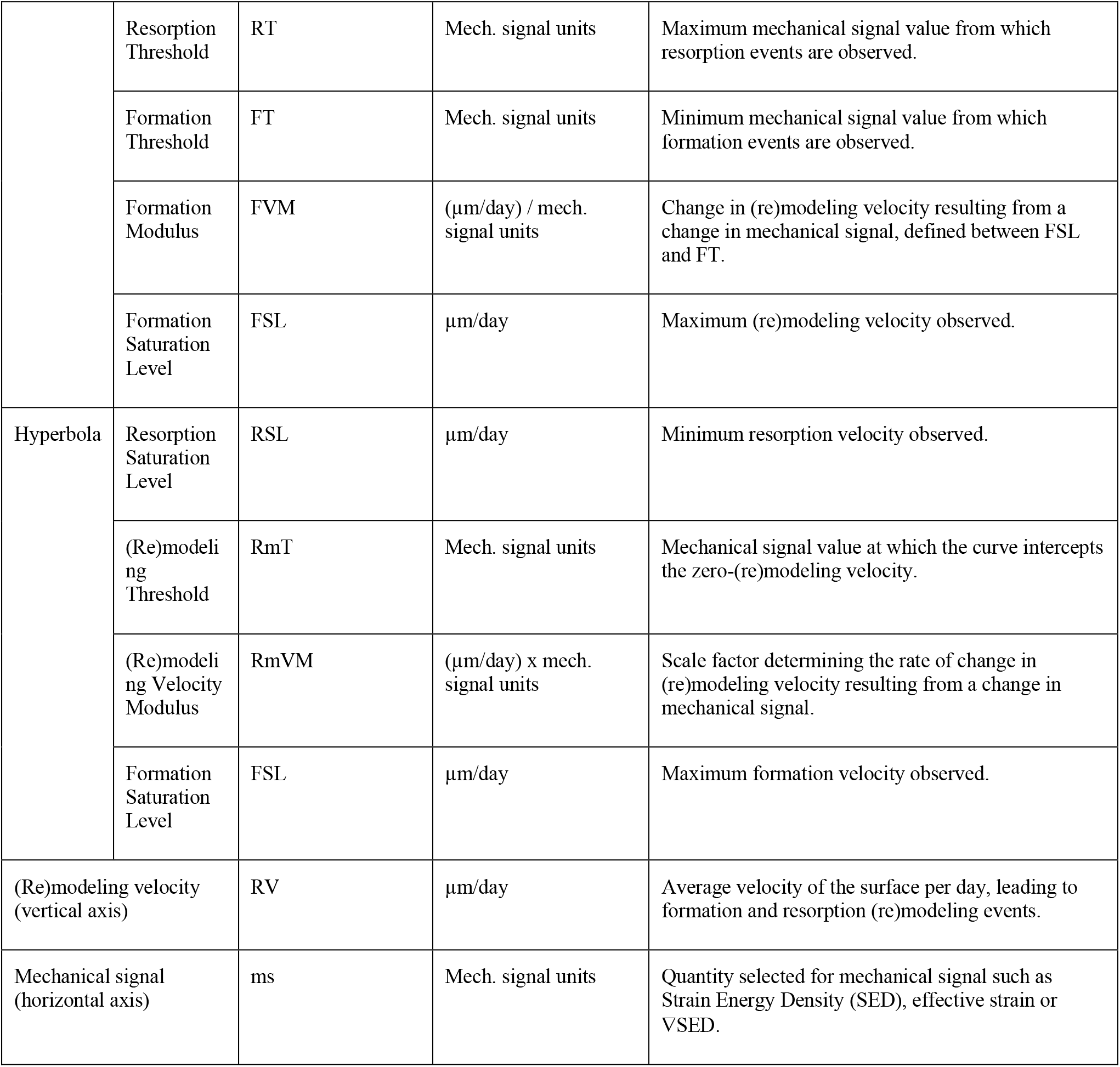
Mechanostat parameters derived from the piecewise linear and hyperbola functions fitted to the (re)modeling velocity curves.

**Figure 1.**
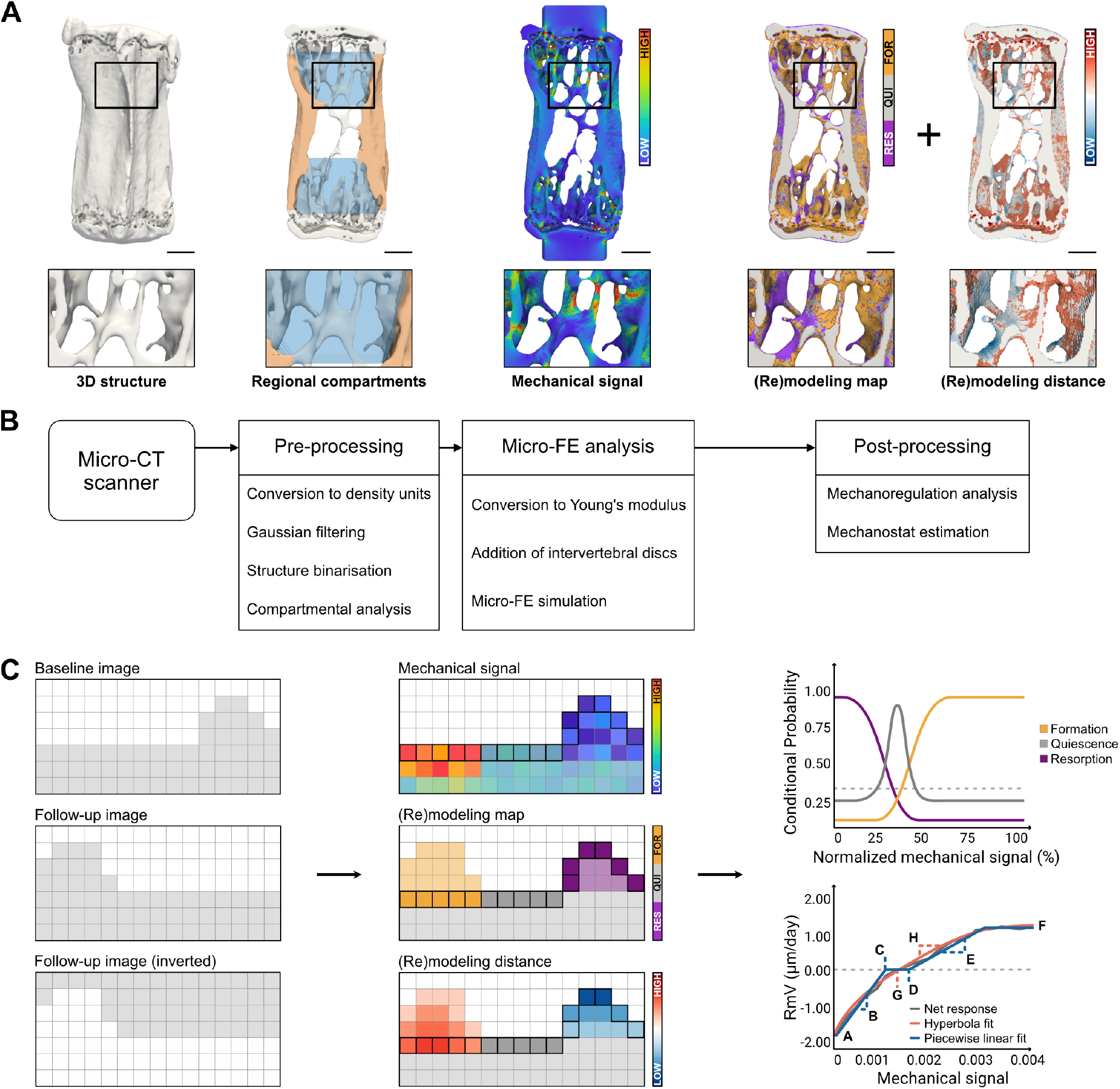
Overview of the computational pipeline for high-throughput analysis of time-lapsed *in vivo* micro-CT mouse caudal vertebra samples. A) Qualitative visualization of a representative sample highlighting the original bone structure, the identification of regional compartments (trabecular compartment in blue and cortical compartment in orange), the local mechanical signal computed as strain energy density (SED) from micro-FE analysis, the (re)modeling map obtained from time-lapsed micro-CT images and the (re)modeling distance associated with surface voxels (Scale bar: 500 µm). B) Diagram of the workflow included in the computational pipeline, from pre-processing micro-CT images to post-processing steps, featuring mechanoregulation analysis. C) Illustration of the workflow for mechanoregulation analysis and estimation of mechanostat parameters: the baseline, follow-up, and inverted follow-up images are considered; for each surface voxel, (re)modeling events are identified by overlapping consecutive time-points, the mechanical signal in the structure is computed with micro-FE and the (re)modeling distance is determined with a distance transform operation (see Materials and Methods). The data is used to compute conditional probability curves for each (re)modeling event (dashed line represents a random probability of occurrence) and a (re)modeling velocity curve (dashed line represents zero (re)modeling velocity), which can be fitted with a piecewise linear function or a continuous hyperbola function to retrieve biologically meaningful parameters. Parameter legend (see Materials and Methods for an extended description): A- Resorption saturation level (RSL), B- Resorption velocity modulus (RVM), C- Resorption threshold (RT), D- Formation threshold (FT), E- Formation velocity modulus (FVM), F- Formation saturation level (FSL), G- (Re)modeling threshold (RmT), H- (Re)modeling velocity modulus (RmVM).

Piecewise linear:

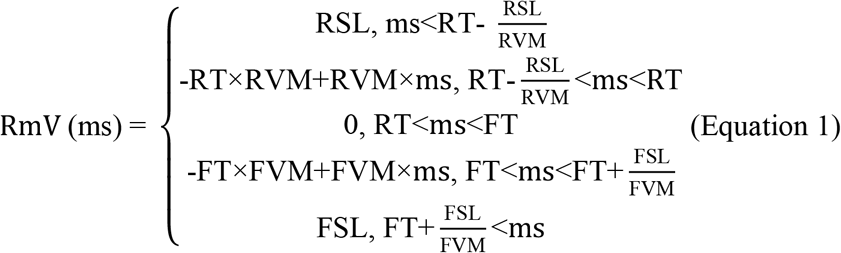

Hyperbola function:

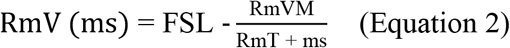

Note that for the hyperbola function, RSL is defined as the value of the RmV function at the minimum mechanical signal value observed.

The quality of the fits was assessed using the root mean squared error (RMSE) between the fitted curve and the corresponding velocity value at each mechanical signal value. The curve fitting was done with Scipy 1.7.3 (Virtanen et al. 2020) and Curve-fit annealing (Reinhardt 2019). Similar to the mechanoregulation analysis using conditional probabilities, the mechanostat (re)modeling velocity curves and associated parameters were estimated for weekly intervals and the 4-week observation period.

We implemented a balanced bootstrapping approach (Dvison et al. 1986) to characterize the distribution of the parameters estimated from the mathematical functions fitted to the (re)modeling velocity curves. Samples were randomly resampled 2500 times per group to generate a synthetic group of the same size, and the corresponding (re)modeling velocity curves were determined and fitted with the piecewise linear and hyperbola functions, yielding the estimations of the mechanostat parameters.

### Frequency dependency of estimated mechanostat parameters

The logarithmic function used previously by Scheuren et al. (2020) was considered here to evaluate the dependency of mechanostat parameters with cyclic loading frequency. Specifically, the median values of the parameter distributions generated with the balanced bootstrapping approach were plotted for each loading frequency and fitted with a logarithmic regression curve (Equation 3). The quality of the fit was assessed with the Pseudo-R^2^ (Schabenberger and Pierce 2001), which enables comparing the quality of fitted relationships for parameters with different magnitudes.

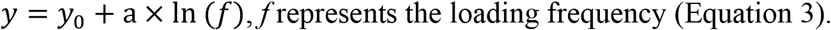

### Statistical analysis

Statistical analysis was performed with Python 3.10.5, using the packages SciPy 1.7.3 (Virtanen et al. 2020) and Scikit_posthocs 0.7.0 (Terpilowski 2019), and in R (R Core Team 2022). Longitudinal measurements of bone structural parameters were also analyzed through repeated measurements ANOVA (Scheuren et al. 2020), implemented as a linear mixed model from the lmerTEST package (Kuznetsova et al. 2017) after inspection of linear regression diagnostics plots. As described by Scheuren et al. (2020), the between-subjects effect was allocated to the groups, while within-subjects effects were allocated to time and time-group interactions. Random effects were allocated to the animal to account for the natural differences in bone morphometry in different mice. In instances where a significant interaction effect (group∗time) was found, a Tukey post-hoc multiple comparisons test was performed. All other parameters were first checked for normality using the Shapiro-Wilk test. Non-normally distributed parameters (CCR values) were presented with their median and inter-quartile range (IQR). Estimated mechanostat parameters were presented with the value of the fitted curve for each case and the IQR of the parameter distribution estimated with the balanced bootstrapping approach. Subsequently, non-parametric tests (Mann-Whitney U-test; Kruskal-Wallis followed by Conover-Iman test for multiple comparisons, corrected by Bonferroni-Holm method) were chosen based on the result of the normality test and indicated accordingly for each comparison in the corresponding figure or table caption. Differences between the conditional probability curves were assessed with Mann-Whitney U-test, for normalized mechanical signal values above 95%. Correlation between (re)modeling velocity curves from different groups was assessed with Pearson Correlation Coefficient. Correlations of static and dynamic morphometry parameters obtained with the IPL and Python compartmental analysis methods (Supplementary Figure 1) and between estimated and ground truth (re)modeling clusters’ volume estimation (Supplementary Figure 3) were determined with Spearman’s correlation coefficient. Significance was set at p < 0.05 in all experiments; otherwise, significance levels are reported.

## Results

### Performance comparison of mechanical signal descriptors for mechanoregulation analysis

First, we extended previous results (Scheuren et al. 2020) by comparing SED, effective strain, and ∇SED and their ability to quantify mechanoregulation information from time-lapsed *in vivo* micro-CT data. Figure 2A illustrates the conditional probability curves for each combination of mechanical signal and group between weeks 0-4. This qualitative evaluation highlighted that, for all mechanical signal descriptors, resorption was confined within a small interval of low mechanical signal values (normalized mechanical signal < 10% for SED and ∇SED and normalized mechanical signal < 29% for effective strain), where it was associated with a higher conditional probability of occurrence.

**Figure 2.**
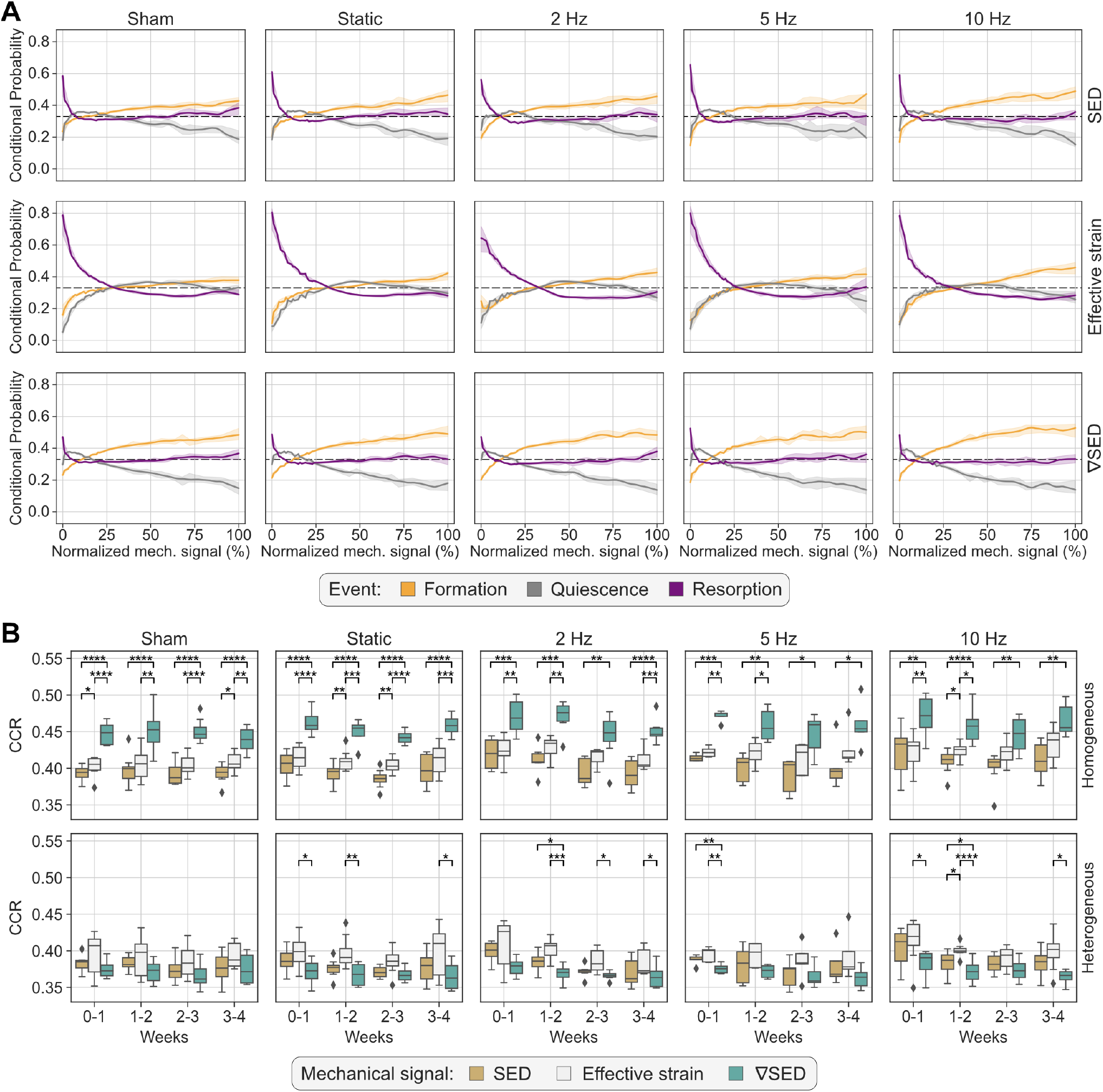
Quantification of mechanoregulation information from time-lapsed *in vivo* micro-CT image data. A) Conditional probability curves connecting the mechanical environment with (re)modeling events, computed for all groups and mechanical signal descriptors considered (SED, effective strain, and ∇SED), using homogeneous material properties. The plots show the mean probability line per group after applying a LOWESS operation for the interval 0-4 weeks and its corresponding 95% confidence interval. The dashed line at 0.33 identifies the probability of a random event for a ternary classification case. B) Comparison of correct classification rate (CCR) values obtained by SED, effective strain, and ∇SED as local mechanical signal descriptors computed from micro-FE with homogeneous or heterogeneous material properties. Higher CCR values indicate higher sensitivity to retrieve mechanoregulation information. Similar to A), the analysis also considers the interval of 0-4 weeks. Statistical significance was determined by the Conover-Iman test, corrected for multiple comparisons by the Holm-Bonferroni method. Statistical significance legend: *p < 0.05, **p < 0.01, ***p < 0.001, ****p < 0.0001.

Furthermore, for higher magnitudes of the mechanical signal, the conditional probability curves displayed a more stochastic pattern oscillating around the random probability of 0.33 for SED and ∇SED and stabilizing below this value for effective strain. ∇SED provided the best discriminative ability for formation and quiescence events for all groups, supported by statistically significant differences when comparing the difference between the conditional probability associated with formation and quiescence between the mechanical signal descriptors (p<0.001 for ∇SED-SED, ∇SED-effective strain, and SED-effective strain). For completeness, these comparisons focused on normalized mechanical signal values above 95%, which comprised a median (IQR) number of voxels per normalized mechanical signal value of 26 (21 – 32) formation voxels, 2 (1 – 3) quiescence voxels and 11 (8 – 16) resorption voxels for SED; 57 (46 – 69) formation voxels, 5 (3 – 7) quiescence voxels and 25 (17 – 35) resorption voxels for effective strain; 27 (22 – 32) formation voxels, 1 (0 – 2) quiescence voxels, and 15 (10 – 19) resorption voxels for ∇SED. These results were also substantiated by the increasing separation between both curves with increased mechanical signal magnitude (Figure 2A). The 10 Hz group achieved the widest difference between these events with 38% for ∇SED, in comparison to 33% for SED and 20% for effective strain, respectively, while the sham-loaded group showed a maximum difference of 33%, 24% and 12% for ∇SED, SED, and effective strain, respectively. Conversely, effective strain showed the best association with resorption events based on the sharp increase in the conditional probability for this event within the interval of low mechanical signal values (normalized mechanical signal < 29%), reaching its maximum conditional probability value between 64% (2 Hz group) and 80% (static and 5 Hz groups), contrasting with values ranging between 47% (sham-loaded group) and 52% (5 Hz group) for ∇SED.

CCR was computed from the conditional probability curves as a proxy of the amount of mechanoregulation information retrieved, quantifying the number of (re)modeling events correctly classified. Our analysis showed that ∇SED consistently achieved the best performance for the micro-FE analysis with homogeneous material properties, followed by effective strain and SED (Figure 2B). Across all groups and all time-points, CCR values for ∇SED were significantly higher than those of SED (Figure 2B). A similar result was observed for ∇SED and effective strain, although no statistical differences were found for the groups with loading frequencies of 5 Hz and 10 Hz after week 2 (Figure 2B). For the interval of 0-4 weeks, CCR values for ∇SED were also significantly higher than those from SED (p<0.001) and effective strain (p<0.05) for all groups except for the 10 Hz group (Table 2). The effect of increasing loading frequencies was also noticeable in the corresponding increase in CCR values in the same period (Table 2). Conversely, for the micro-FE analysis with heterogeneous material properties, effective strain showed the best association with (re)modeling events, followed by SED and ∇SED, albeit the accuracy was lower than what was observed for micro-FE with homogeneous material properties, both in the weekly analysis (Figure 2B) and between 0-4 weeks (Table 2). Furthermore, there was a weaker increasing trend of CCR values with increasing loading frequencies (Table 2) in comparison to the micro-FE analysis with homogeneous material properties.

**Table 2.** Correct classification rate (CCR) for all groups, mechanical signal descriptors, and material properties analyzed for weeks 0-4. Data presented as “median (IQR)”. Statistical significance legend: a – “SED – Effective strain”, b – “Effective strain – ∇SED”, c – “SED – ∇SED”; ns: not significant, *p < 0.05, **p < 0.01, ***p < 0.001, ****p < 0.0001. Statistical significance was determined with Conover’s test corrected for multiple comparisons with a step-down method using Bonferroni-Holm adjustments.

| Material properties | Group | Mechanical signal |  |  | p-value |
| --- | --- | --- | --- | --- | --- |
| | | SED | Effective strain | $\nabla\text{SED}$ | |
| Homogeneous | Sham | 0.378 (0.374 - 0.391) | 0.400 (0.392 - 0.404) | 0.421 (0.413 - 0.440) | *a, **b, ****c |
|  | Static | 0.389 (0.375 - 0.395) | 0.405 (0.403 - 0.410) | 0.434 (0.427 - 0.448) | *b, ***c |
|  | 2 Hz | 0.402 (0.387 - 0.404) | 0.411 (0.407 - 0.417) | 0.450 (0.430 - 0.451) | *b, ***c |
|  | 5 Hz | 0.396 (0.387 - 0.397) | 0.404 (0.399 - 0.422) | 0.446 (0.439 - 0.454) | *a, *b, ***c |
|  | 10 Hz | 0.417 (0.382 - 0.443) | 0.418 (0.395 - 0.428) | 0.461 (0.431 - 0.489) | ns |
| Heterogeneous | Sham | 0.381 (0.373 - 0.387) | 0.423 (0.410 - 0.432) | 0.371 (0.366 - 0.374) | *b, **c |
|  | Static | 0.386 (0.379 - 0.392) | 0.423 (0.416 - 0.434) | 0.377 (0.364 - 0.381) | *b, **c |
|  | 2 Hz | 0.387 (0.387 - 0.397) | 0.430 (0.399 - 0.434) | 0.382 (0.375 - 0.391) | *b, **c |
|  | 5 Hz | 0.394 (0.387 - 0.398) | 0.420 (0.412 - 0.431) | 0.380 (0.372 - 0.391) | ns |
|  | 10 Hz | 0.397 (0.387 - 0.435) | 0.427 (0.414 - 0.447) | 0.395 (0.365 - 0.408) | ns |

### Quantification of (re)modeling velocity curves and mechanostat parameters

Next, we investigated bone mechanoregulation from time-lapsed *in vivo* micro-CT data by deriving (re)modeling velocity curves and fitting piecewise linear and hyperbola functions to estimate the corresponding mechanostat parameters, namely formation and resorption saturation levels (FSL and RSL), velocity modulus (FVM and RVM) and thresholds (FT and RT) for the piecewise linear function and FSL and RSL, (re)modeling velocity modulus (RmVM) and (re)modeling thresholds (RmT) for the hyperbola function, respectively. We analyzed consecutive pairs of scans (Figure 3, Supplementary Table 1) one week apart and summarized the group differences over the four-week period of the study (Table 3).

**Table 3.**
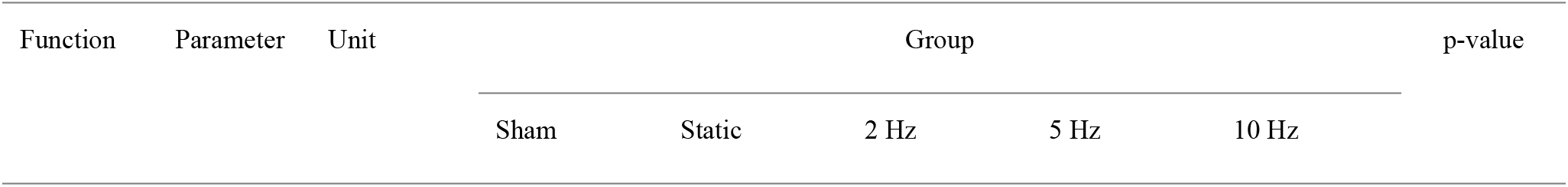
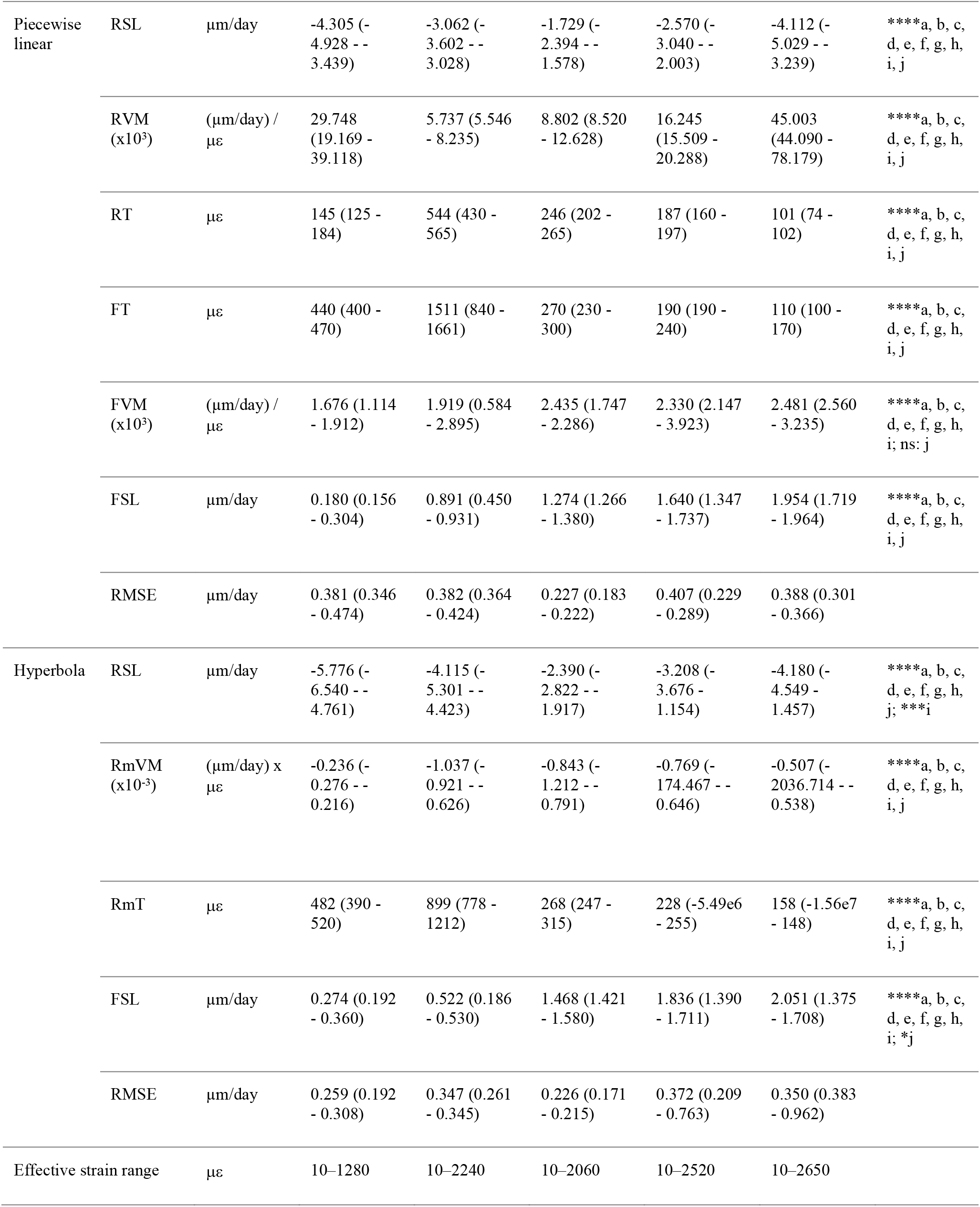
Parameters of the mathematical functions fitted to the estimated mechanostat group average (re)modeling velocity curves for the interval weeks 0-4. Data presented as “parameter (IQR)”, where the IQR was estimated using the balanced bootstrapping approach described in the methods (see section “Mechanostat (re)modeling velocity curve and parameter derivation”) to provide an estimate for the variability of each parameter. Root mean squared error (RMSE) was used to characterize the quality of the fit. The row “Effective strain range” indicates the range of mechanical signal values from which the fit of the mathematical functions was derived. Mechanical signal values obtained from micro-FE analysis using homogeneous material properties. Parameter legend (see Materials and Methods and Table 1 for an extended description): Resorption saturation level (RSL), Resorption velocity modulus (RVM), Resorption threshold (RT), Formation threshold (FT), Formation velocity modulus (FVM), Formation saturation level (FSL), (Re)modeling threshold (RmT), (Re)modeling velocity modulus (RmVM). Statistical significance legend: a – “Sham – Static”, b – “Sham – 2 Hz”, c – “Sham – 5 Hz”, d – “Sham – 10 Hz”, e – “Static – 2 Hz”, f – “Static – 5 Hz”, g – “Static – 10 Hz”, h – “2 Hz – 5 Hz”, i – “2 Hz – 10 Hz”, j – “5 Hz – 10 Hz”; ns: not significant, *p < 0.05, **p < 0.01, ***p < 0.001, ****p < 0.0001. Statistical significance was determined with Conover’s test corrected for multiple comparisons with a step-down method using Bonferroni-Holm adjustments.

**Figure 3.**
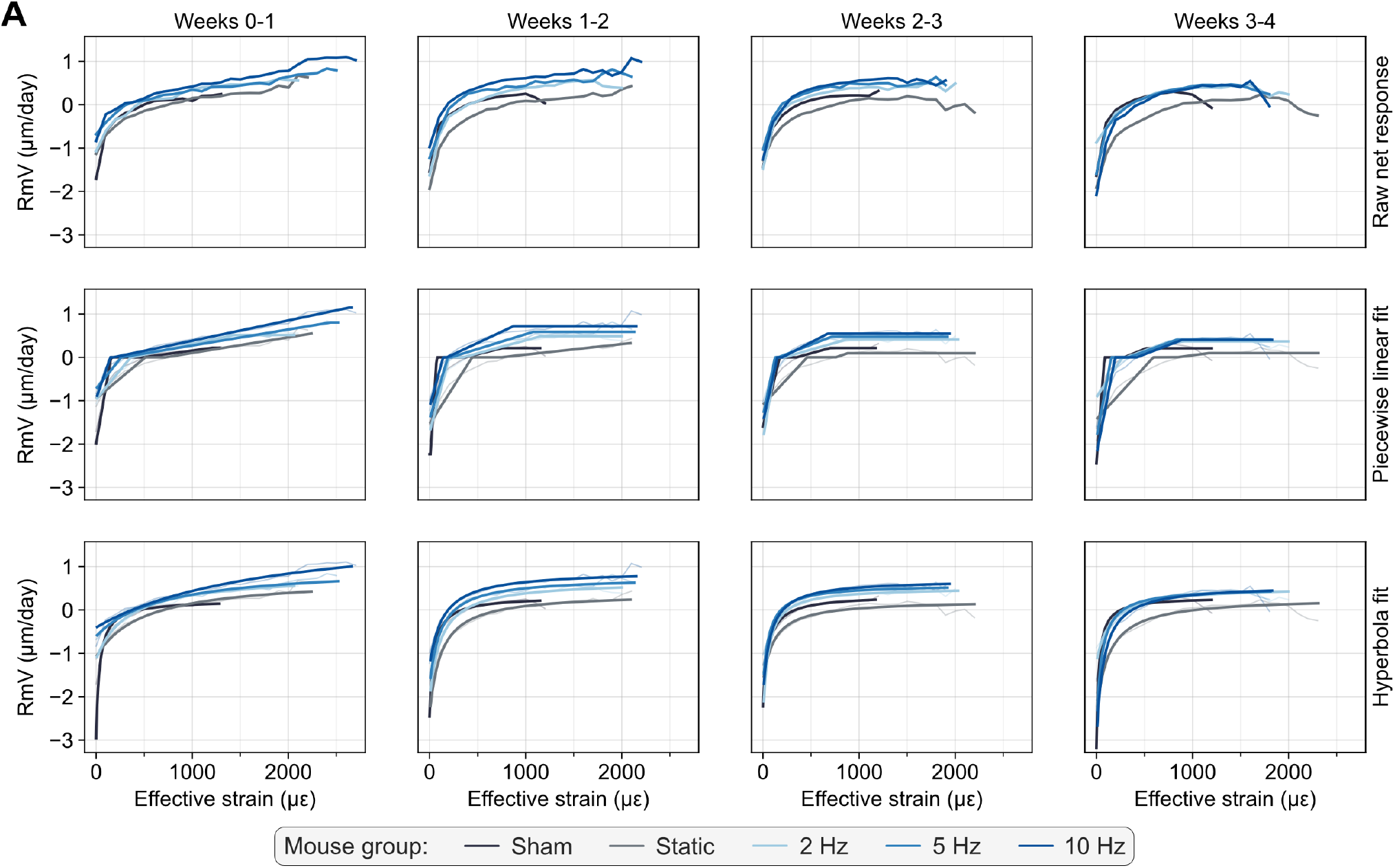
Estimation of the mechanostat (re)modeling velocity (RmV) curve from time-lapsed *in vivo* micro-CT imaging data, illustrated with the average raw net response (top row) per group, the fitted piecewise linear functions (middle row), as described in the mechanostat theory and continuous hyperbola functions (bottom row). Data points are filtered such that at least three mice are averaged for each mechanical signal value. Mechanical signal values obtained from micro-FE analysis using homogeneous material properties.

The raw net response curves shown in Figure 3 resemble the shape of the mechanostat schematic proposed by Frost (1987) from the disuse to the adapted and mild overload windows, providing a qualitative validation of the output, which evaluated supraphysiological cyclic loads applied to the mouse vertebrae. Furthermore, increasing loading frequency led to a linear translation of the derived curves towards higher RmV values, with formation events starting from lower mechanical signal threshold values (Figure 3). Quantitatively, the parameters derived from the fitted curves showed a significant increase in the FSL and decreased RT and RmT values with increasing loading frequency for both the weekly and the 0-4 weeks analysis (Table 3, Supplementary Table 1). Conversely, the RSL values decreased significantly for increasing loading frequencies (Supplementary Table 1), especially for weeks 3-4. Reinforcing the effects of different loading frequencies on bone adaptation, statistically significant differences were observed between estimated parameters between groups and time-points (Supplementary Figures 3-6).

The RmV curves also allowed characterizing time-lapsed bone adaptation for each group, where the range of RmV values decreased weekly (Figure 3) and converged towards comparable values between groups. Overall, this result indicated that the magnitude of the mechanical signal in weeks 3 and 4 no longer induced the same strong (re)modeling responses observed in the first two weeks.

FSL values followed a similar pattern for each group over the four weeks (Supplementary Table 1). In the first week, the RmV curves highlighted the acute response to supraphysiological loading since most loaded groups did not reach a plateau at their FSL value, which only occurred in subsequent time-points. This progression was visible in the raw net response curves and the fitted mathematical functions (Figure 3). Concurrently, RVM and FVM values (Supplementary Table 1) increased significantly over time such that, at weeks 0-1, only regions of either high or low effective strain could elicit the strongest response associated with the estimated formation and resorption saturation levels, respectively. Over time, as the bone structure adapted to supraphysiological loading, the extent of regions of either high or low effective strain decreased, also visible in the decrease in the range of mechanical signal values (except for the sham-loaded group), and FSL were reached for lower mechanical signal values (Figure 3, Supplementary Table 1, Supplementary Figures 3-6). The RmV curves showed that even groups loaded with higher frequencies (5 Hz and 10 Hz) evolved from a state of predominant formation in the first two weeks towards physiological (re)modeling conditions, where resorption was tightly regulated within an interval of low mechanical signals and in agreement with the conditional probability curves shown previously (Figure 2). These observations were also corroborated by increased Pearson correlation coefficients between (re)modeling velocity curves of 5 Hz and 10 Hz loaded groups and the sham-loaded group.

Specifically, these increased from 0.828 and 0.822 (p<0.0001) for weeks 0-1 to 0.908 and 0.883 (p<0.0001) in weeks 3-4, for the 5 Hz and 10 Hz groups, respectively, supporting comparable bone (re)modeling responses between these groups except for their mechanosensitivity, as seen in the differences between FT and RT values. Comparably, the RmV curves between loaded groups also evolved towards a more uniform bone (re)modeling response based on the increase in the Pearson correlation coefficients between the 2 Hz and 10 Hz groups and the 5 Hz and 10 Hz, from 0.872 and 0.874 to 0.899 and 0.925, respectively.

Regarding the effects of static and dynamic loads, we observed that static loading still induced an anabolic response in the first week, characterized by a higher FSL than the sham-loaded group (Figure 3). However, in weeks 1-2 and 2-3, the static group already matched the FSL values of the sham-loaded group suggesting a return to a physiological (re)modeling condition, and eventually reached a lower FSL in weeks 3-4. RT and FT were significantly higher in the static group than in the sham-loaded group across all weeks, indicating that this loading condition still produced high strains in the structure. Likewise, RmV curves also reflected net bone volume changes by integrating them with respect to time and the mechanical signal. While the sham and static groups showed a negative volume change over four weeks, cyclically loaded groups showed increasing net positive changes with increasing loading frequency for the same interval.

Comparing the piecewise linear and hyperbola functions, the RMSE of the fitted curves indicated that these could be determined reliably and accurately, with an average RMSE of 0.357 and 0.314 µm/day for the former and latter, respectively, over the 4-week interval (Table 3) and yielding even lower RMSE values for the weekly analysis (Supplementary Table 1), partially given the lower magnitude of (re)modeling velocities observed. Additionally, lower RMSE values coupled with a wider range of (re)modeling velocities obtained with the fitting of the hyperbola function suggested more accurate curve fits than with the piecewise linear function, which is especially crucial for the quantification of (re)modeling thresholds in the region where the (re)modeling velocity is zero (Figure 4A, Supplementary Table 1).

**Figure 4.**
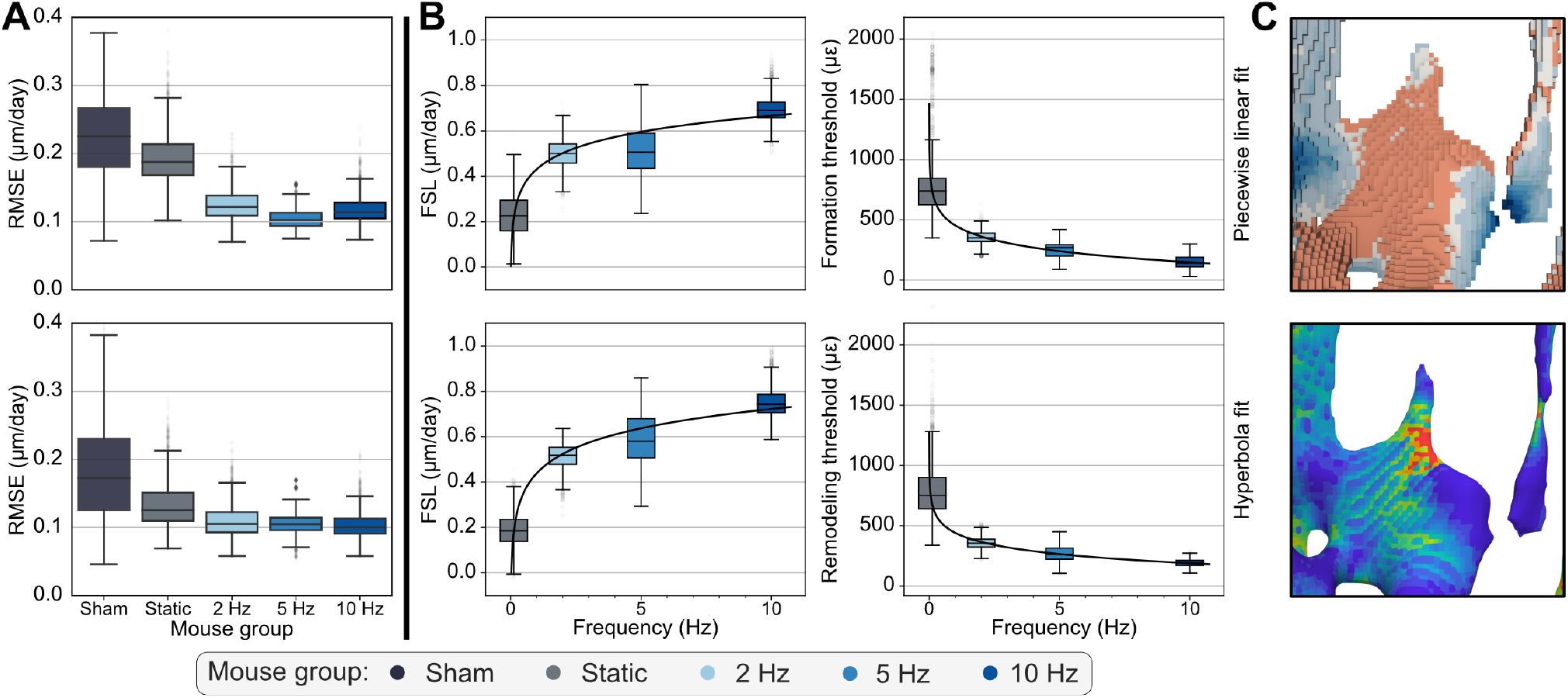
A) Root mean squared errors (RMSE) associated with the piecewise linear (top row) and hyperbola (bottom row) fitted functions for the analysis of weeks 1-2 are shown, highlighting that the hyperbola function consistently achieved lower errors than the piecewise linear function. B) Logarithmic relationships fitted to the bootstrapped distributions of mechanostat parameters estimated from the piecewise linear (top row) and hyperbola (bottom row) functions fitted to the (re)modeling velocity curves for the weeks 1-2. Formation saturation levels (FSL), formation, and (re)modeling thresholds (FT and RmT) were among the parameters that followed a logarithmic trend throughout the four weeks of the study. C) Qualitative visualization linking (re)modeling distance measurements with the mechanical environments *in vivo* as effective strain.

Finally, our analysis also investigated the trend followed by the mechanostat parameters derived from the fitted (re)modeling velocity curves with loading frequency. A logarithmic function was suitable for several parameters estimated from the piecewise linear and hyperbolic fits (Figure 4B, Supplementary Table 2). For the piecewise linear function, FSL was accurately modeled by a logarithmic function across all time-points (Supplementary Table 2, Figure 4B for the interval 1-2), especially from week 1 onwards and for the analysis over the 4-week observational period (weeks 0-4). FT values also followed the same trend for weeks 0-1 and 1-2, while RT showed the same behavior for all weekly time-points between weeks 1 and 4 (Supplementary Table 2). Similarly, for the hyperbola function, FSL was also accurately modeled by a logarithmic function across all weekly time-points and between weeks 0-4 (Supplementary Table 2, Figure 4B for the interval 1-2). Besides, RmT followed a logarithmic relationship until week 2, analogous to the FT estimated with the piecewise linear function.

## Discussion

The present study evaluated mechanoregulation in trabecular bone adaptation and quantitatively characterized the effects of loading frequencies on bone adaptation using a novel method to estimate (re)modeling velocity curves and their mechanostat parameters from time-lapsed *in vivo* mouse vertebra micro-CT data. Crucially, we showed that such RmV curves could be accurately determined and that several parameters obtained from them followed a logarithmic relationship with loading frequency, further supporting the trend observed previously for the change in bone volume fraction over the 4-week observation period (Scheuren et al. 2020).

First, we consolidated key factors that yield the best association between mechanical stimuli and local bone (re)modeling using existing methods for mechanoregulation analysis. Our results revealed that micro-FE analysis with homogeneous material properties achieved the best performance in recovering mechanoregulation information. These homogeneous material properties were determined from *in vivo* experiments using strain gauges attached to the cortical walls of the murine vertebrae, such that the strains computed with micro-FE would correlate best with the *in vivo* measurements (Webster et al. 2008). Conversely, the mathematical relationship relating bone mineral density and Young’s Modulus for the micro-FE analysis with heterogeneous material properties was derived from nanoindentation experiments on the mandibular condyles of pigs (Mulder et al. 2007). Given the differences in animal models and anatomical sites, it is plausible that this relationship could influence the accuracy of the Young’s Modulus conversion and, ultimately, the strain distribution obtained from micro-FE with heterogeneous material properties. Indeed, we observed that this relationship led to a higher average Young’s Modulus value than the value used for homogeneous material properties (Supplementary Material 1.2). In alignment with our results, previous work on the murine tibia (Oliviero et al. 2021) has shown that micro-FE analysis with homogeneous material properties achieved the highest correlation between experimental and estimated material properties, while micro-FE with heterogeneous material properties of the lumbar vertebra L6 did not improve the prediction of failure force in comparison to homogeneous material properties (Harris et al. 2020). In any case, other applications where more significant changes in bone mineralization are expected, such as during fracture healing of cortical bone in the mouse femur (Tourolle né Betts et al. 2020), were more accurately modeled with heterogeneous material properties. Still, in the context of load-induced trabecular bone adaptation as explored in this work, the micro-CT images did not show large dynamic ranges in bone mineralization and changes in bone volume (Lambers et al. 2011, Oliviero et al. 2021), suggesting that the use of homogeneous material properties in micro-FE analysis is appropriate to model mechanically driven bone adaptation.

From a mathematical modeling perspective, the mechanostat theory (Frost 1987) is an established paradigm to describe bone adaptation in response to mechanical loading that has also been successfully applied in preclinical *in silico* models (Levchuk et al. 2014, Pereira et al. 2015, San Cheong et al. 2020, San Cheong et al. 2020). The analysis proposed in this work enables a direct estimation of such a relationship from time-lapsed *in vivo* micro-CT data and can be applied in a sample-specific or group-wise fashion and for an arbitrary time interval between the input images. We also proposed a nomenclature of mechanostat parameters, unifying the descriptions used in previous studies. For instance, the change in bone material in response to mechanical loading was originally named bone turnover and bone growth by Frost (1987) and later adapted to growth velocity using a detailed mathematical framework *in silico* (Levchuk et al. 2014, Goda et al. 2016, Louna et al. 2019). In agreement with these studies and with potential relevance for future *in silico* models, we chose to also focus our analysis on velocity rather than change in bone mass, spatially resolving the displacement of surface voxels between time-points. In fact, recent advances in the context of *in silico* single-cell mechanomics (Boaretti et al. 2023) have associated positive and negative velocities with the activity of osteoblasts and osteoclasts, respectively. Still, converting the output to other units of interest, including a change in bone volume as originally proposed, is equally possible. Furthermore, we opted for (re)modeling velocity as it fits the context of bone adaptation where there can be negative and positive growth (Hadjidakis and Androulakis 2006). Conversely, formation/resorption thresholds and saturation levels were consistent with previous approaches (Levchuk et al. 2014, San Cheong et al. 2020). Regarding the change of RmV with mechanical signal, it was appropriate to align this term with the naming structure of the remaining parameters and provide an intuitive, succinct description integrating a modulus terminology: formation/resorption/(re)modeling velocity modulus. In this way, we aimed to strengthen the association of these terms with the mechanical signals, as modulus is inherently linked with other mechanical terms that relate a change in a quantity with a change in mechanical strain, such as Young’s modulus, describing the relationship between stress and strain for a given material.

Focusing on the effects of mechanical loading, we observed statistically different FT, RT, and RmT per group. These observations align with the original predictions by Frost (1987), who argued that disease states (leading to long periods of physical inactivity) or increased mechanical demands could increase or decrease these parameters, respectively. In our approach, at the micrometer scale, these thresholds reflect the transition between an interval of lower strains where resorption is predominantly observed and an interval of higher strains where formation is, on average, more frequent and with higher magnitude. As cyclic loading groups with increasing frequency showed more formation events, (re)modeling velocity curves were shifted towards higher velocity values, yielding progressively lower thresholds. Conversely, sham and static groups showed a slight bone loss in the observation period, concordant with higher thresholds. Focusing on mild-overload examples, Rubin and Lanyon (1985) had already noted that the thresholds for bone formation would vary depending on the loading characteristics. Although Frost (1987) could only anticipate the existence of such thresholds, technological advances have enabled identifying possible biological agents that realize them at the cellular level. For instance, the absence of Connexin 43 (Cx43) membrane protein in osteocytes enhanced the responsiveness to mechanical force in mice (Bivi et al. 2013). Similarly, the activation of the Wnt/β-catenin pathway increased the sensitivity of osteoblasts and osteocytes to mechanical loading (Gerbaix et al. 2015, Gerbaix et al. 2021). Conversely, high-magnitude mechanical stress was observed to inhibit canonical Wnt signaling-induced osteoblastic differentiation (Song et al. 2017). These examples highlight adjustable responses to external mechanical loading events by cells implicated in bone (re)modeling events.

On a similar note, Skerry (2006) stated that different loading conditions, such as those induced *in vivo* through varying loading frequencies, produce deviations from the habitual strain stimuli of the structure. Furthermore, he argued that different anatomical sites have specific “customary strain stimulus (CSS)” values to which the structure adapts. Our results align with these beliefs, where different loading frequencies produced significantly different responses (Scheuren et al. 2020), and the RmV curves evolved towards a state where (re)modeling thresholds were very close, suggesting a return to the habitual mechanostat rule and its local CSS value. For this reason, it is understandable that FT and RmT were no longer logarithmic dependent on loading frequency at week 4. Conversely, RT conserved a logarithmic trend with loading frequency for all weeks, which aligns with the tight regulation of resorption events observed in conditional probability and RmV curves. Crucially, an accurate derivation of the mechanostat curve required calibrating the volume estimated for each (re)modeling cluster (Supplementary Figure 3A). For instance, the volume of smaller clusters with a high surface-to-volume ratio was expectedly overestimated by the distance transform operation. This artifact is particularly noticeable for formation clusters where the identification of the neighboring surface from which they emerged required a morphological dilation operation, leading to an increase in the number of surface voxels related to this event. While previous studies (Schulte et al. 2013, Razi et al. 2015, Scheuren et al. 2020) assessing bone mechanoregulation focused exclusively on conditional probabilities, which only consider the frequency of mechanical signal values per (re)modeling event, this volume correction becomes of significant importance in our proposed method, where a new axis focusing on the (re)modeling velocity at each voxel is considered. Ultimately, this observation sustains two key aspects. First, in agreement with previous studies (Lanyon and Rubin 1984, Turner et al. 1995, Robling et al. 2001) that showed that dynamic but not static loads induce an adaptive response, also our estimated RmV curves conserve this hallmark previously observed in the dataset analyzed here (Scheuren et al. 2020). Second, an accurate volume estimation enables the identification of critical setpoints, such as formation and resorption thresholds, where the RmV curve approaches zero. Notably, the interval defined by these thresholds is typically described as a lazy zone, i.e., a range of strains where bone formation and resorption balance each other, analogous to the adapted window proposed by Frost (1987).

In this regard, and in agreement with previous findings in preclinical mouse (Sugiyama et al. 2012, Schulte et al. 2013, Razi et al. 2015, San Cheong et al. 2020) and clinical (Christen et al. 2014) data, our results provided no evidence of the existence of a lazy zone. This was further supported by the lower RMSE values associated with the fitted hyperbola mathematical functions which, by definition, cannot accommodate such an interval. Regardless, the estimated (re)modeling velocity curves agree with previous publications (Schulte et al. 2013, Razi et al. 2015), where resorption seems to be more tightly regulated than formation, based on the width of the interval of mechanical signal values allocated to each (re)modeling event, consistent for all groups and loading frequencies. Furthermore, the estimated (re)modeling rates, ranging between 0 and 3 µm/day, agree with the corresponding mineral apposition and resorption rates previously reported for this dataset (Scheuren et al. 2020) at around 2 µm/day, averaged across all (re)modeling clusters identified. Besides, the decreasing RSL values for increasing loading frequencies observed (Supplementary Table 1), especially for weeks 3-4, also align with previous work on the mouse tibia that showed an increase in the depth of resorption cavities with loading (Birkhold et al. 2017). Given the correction in the curve estimation that ensures accurate volumes, established dynamic morphometry indices characterizing bone formation and resorption rates in a single value can now be expanded into a range of mechanical signals.

On a different note, previous studies (Rubin and Lanyon 1985, Turner et al. 1994) identified a strain threshold of around 1000 µε from which formation events were observed. This estimate seemingly contradicts the conditional probabilities and (re)modeling velocity curves presented (Figure 2A), where formation had a non-zero probability of being observed over the entire range of mechanical signals and positive velocity values started from 200 µε, respectively. However, those studies focused primarily on the effects of the supraphysiological loading applied to the animal models, offsetting effects resulting from physiological activity (e.g., by subtracting reference measurements from the control leg to the loaded leg and defining relative measures). Conversely, our analysis considered all (re)modeling events observed, that is, from both modeling and remodeling processes occurring concurrently. In agreement with previous studies (Burr 2002) describing other non-deterministic processes driving formation and resorption events, it is reasonable that these can occur in locations that a purely mechanically driven system would not favor. Indeed, Lanyon et al. (1982) had already observed bone formation in a sheep model in regions with “functional strains lower than normal values. Furthermore, previously reported values refer to peak strains determined on the external cortical wall of each bone, while our analysis considers local effective strain inside the trabecular compartment, making a direct comparison of magnitudes challenging. Still, notably, Webster et al. (2008) originally reported peak strains measured with strain gauges on the external cortical wall of caudal vertebrae of the same mouse model considered in our work within the same range as those obtained in previous studies.

Nonetheless, there are some limitations to consider in this study. First, although the estimation of RmV curves can be determined in a sample-specific fashion, we observed that the analysis of group average curves was more reliable. These naturally contained more data points which were also filtered such that at least three samples were considered per mechanical signal value. Eventually, these factors were vital to producing relatively smooth RmV curves and enabling consistent and plausible piecewise linear, and hyperbola fits. In any case, as previous work has focused on group average results both *in vivo* (Schulte et al. 2013, Razi et al. 2015, San Cheong et al. 2020) and *in silico* (Levchuk et al. 2014, San Cheong et al. 2020, Boaretti et al. 2023), our analysis still aligns with such standard practices. Second, contrasting with conditional probability-based approaches, (re)modeling events are no longer characterized separately since our approach yields a single curve representing the average RmV describing the net effect of (re)modeling events for a given mechanical signal. Effectively, this feature prevents our method from capturing more subtle trends observed in the conditional probability curves (e.g., the slight increase in the conditional probability of resorption for higher mechanical signal values). Nonetheless, our goal was to derive a relationship in alignment with the mechanostat theory which, by definition, also does not describe (re)modeling events independently. Although conditional probability curves showed that these events could occur across the entire range of mechanical signals and highlight the interrelated effect of mechanical and biological cues governing targeted and non-targeted bone (re)modeling (Burr 2002, Parfitt 2002, Schulte et al. 2013), we consider our approach complementary to this probability-based method. Still, as different mechanical signal quantities performed differently for formation and resorption events (Figure 2A), future approaches can attempt to combine both methods and derive separate RmV curves for formation and resorption using the mechanical signal that best associates with each event.

Likewise, our approach was not designed to investigate the spatial and temporal links between surface (re)modeling events (i.e., analysis of the spatial/temporal distribution of formation, quiescent and resorption regions), analogous to previously proposed methods (Birkhold et al. 2015). Indeed, in the mouse bone and with the resolution of the micro-CT used here, one cannot distinguish between pure remodeling and modeling events, which we collectively refer to as (re)modeling (Huiskes et al. 2000, Schulte et al. 2013). Therefore, it is expected that physiological (re)modeling also took place during the 4-week observation period. One can argue that such events are continuously happening and would not be as mechanically driven, yielding flatter (re)modeling velocity curves. Accordingly, our observations of the sham group, where (re)modeling events should be most noticeable, feature lower FSL, FVM, and RmVM values while conserving a tighter regulation of resorption events. With potential applications for clinical data focusing on disease states, this observation would be particularly impactful as current *in vivo* clinical imaging modalities are still affected by noise and motion artifacts, such that reliably identifying (re)modeling events is challenging (Christen et al. 2018). Eventually, longer intervals between scans would be required to produce time-lapsed data with sufficient changes that could yield RmV curves depicting a more explicit relationship with mechanical signals computed from micro-FE.

It should be noted that the current micro-CT image resolution also challenges an accurate identification of sub-voxel phenomena. Indeed, such information would help to elucidate the assumption considered in our (re)modeling velocity estimation that the (re)modeling distance measured for each voxel surrounding a (re)modeling cluster can be linearly scaled to match the calculated volume of the cluster. For the same reason, this factor also implies that the proposed method cannot recover single-cell behavior. Nonetheless, loading frequency was positively correlated with the number of osteocytes recruited in response to an increase in applied strain (Lewis et al. 2017), with a particular focus on bone formation in a murine metatarsal model. Additionally, in a rat tibia model, increasing loading frequency was associated with a decrease in the estimated peak microstrain triggering periosteal bone formation and an increase in the rate of bone formation per microstrain (Hsieh and Turner 2001). Combined, these results would emerge as an increase in FVM and a decrease in FT values with increasing frequency, which is what our RmV curves show until week 3. Furthermore, the decreased anabolic response observed in the RmV curves for high strains may also be linked to a decrease in mechanosensitivity resulting from increased cell stiffness, as previously reported for such high strain values (Nawaz et al. 2012). Therefore, the trends estimated with the mechanostat (re)modeling velocity curves could be leveraged by *in silico* simulations that also rely on time-lapsed *in vivo* micro-CT data as input, such as novel agent-based models that simulate individual cell populations in 3D (Tourolle 2019, Boaretti et al. 2023) and, with that, improve the accuracy of their predictions with respect to *in vivo* data. For this reason, we chose to express RmV curves using effective strain, as this signal has already been implemented successfully in such *in silico* models. Still, our approach is compatible with any voxel-based mechanical signal descriptor, including ∇SED which showed the best association with (re)modeling events (Figure 2) or other signals implemented in *in silico* models of trabecular bone remodeling. Furthermore, our results demonstrating that several parameters estimated from the mechanostat also follow a logarithmic relationship with loading frequency can help to calibrate such models and investigate loading frequency-dependent responses *in silico*. With the advent of more powerful imaging methods, cell populations may soon be efficiently measured from *in vivo* samples and compared with the results of these *in silico* models. Additionally, our approach can support preclinical *in vivo* studies focusing on bone mechanoregulation. Previous work exploring the effects of aging and degenerative conditions described changes in conditional probabilities between young and aged groups (Razi et al. 2015), while studies focusing on pharmaceutical interventions characterized changes in global morphometry indices and micro-FE properties (Roberts et al. 2020), and also reported synergistic effects between specific treatments and mechanical loading in a mouse model of osteoporosis (Levchuk et al. 2014). Therefore, as Frost (1987) had already anticipated the effects of medications on the thresholds bounding the adapted window, our approach provides a quantitative framework to support this analysis. Interventions leading to changes in (re)modeling thresholds (FT, RT, RmT) could suggest changes in the sensitivity to loading while varying (re)modeling modulus (FVM, RVM, RmVM) would relate to changes in the magnitude of the response. Ultimately, it could be possible to identify which intervention would help to counter such disruptions in bone mechanoregulation that follow degenerative conditions.

In conclusion, we have presented a novel method to estimate (re)modeling velocity curves and their parameters from time-lapsed *in vivo* micro-CT data. Furthermore, we applied this approach to evaluate the effects of different loading frequencies on time-lapsed changes in bone microarchitecture by quantifying critical parameters describing bone mechanoregulation, such as formation saturation levels and (re)modeling thresholds. Crucially, we reinforced previous results that revealed a logarithmic relationship of bone volume change with loading frequency by showing that mechanostat parameters estimated from RmV curves, such as (re)modeling thresholds and formation and resorption saturation levels, are also logarithmically dependent on loading frequency. Altogether, we expect these results to support future *in silico* and *in vivo* studies comparing the effects of mechanical loading and pharmaceutical treatment interventions on bone mechanoregulation and bone adaptation and, ultimately, identify more effective treatment plans that can be translated into clinical settings.

## Author Contributions

**Study design**: FM, AS, RM. **Study conduct**: FM, DB, MW. **Data collection**: FM. **Data analysis**: FM. **Data interpretation**: FM, DB, FS, RM. **Drafting manuscript**: FM. **Revising manuscript content**: FM, DB, MW, AS, FS, RM. **Approving final version of manuscript**: FM, DB, MW, AS, FS, RM. RM takes responsibility for the integrity of the data analysis.

## Acknowledgments

This manuscript was released as a preprint at bioRxiv (C. Marques et al. 2023).

## 1 Data Availability Statement

Data and code will be made available upon reasonable request.

