## Supplementary Material for "Mechanostat parameters estimated from time-lapsed *in vivo* micro-computed tomography data of mechanically driven bone adaptation are logarithmically dependent on loading frequency"

frontiers

### *Supplementary Material*

#### 1 Supplementary Material 1

##### 1.1 High-performance computing analysis pipeline

The analysis performed in this work involved a substantial use of computational resources. For this reason, we have implemented an automated pipeline, compatible with modern programming languages and high-performance computing (HPC) platforms, that can dramatically expedite the throughput of the analysis for such large-scale *in vivo* studies and support subsequent *in silico* modeling of biological processes.

###### 1.1.1 Automated compartmental analysis of the murine caudal vertebrae

The first stage concerns the identification of regional compartments inside the murine vertebrae, namely the cortical and trabecular regions. Our proposed Python implementation was validated against the established IPL version to enable its use in all subsequent steps. The reproducibility of the analysis was implicitly evaluated since all operations in the pipeline are deterministic, and the pipeline ran successfully for all samples. Dice Similarity Coefficient (Dice 1945, Sørensen 1948) was quantified to describe the accuracy of the compartments generated with the proposed Python pipeline compared to the IPL implementation. Static and dynamic morphometric parameters were computed for the trabecular compartments generated with each method using a previously established direct 3D approach (Hildebrand et al. 1999) in IPL and compared using Spearman correlation coefficient (Kokoska and Zwillinger 2000, Virtanen et al. 2020) to characterize the accuracy and sensitivity of the proposed implementation to extract biological information about the bone microarchitecture. In addition, a group-wise weekly comparison of morphometric parameters obtained with each pipeline was performed to assess the sensitivity of the Python version to quantify microarchitectural differences in the data.

The images from all time-points were successfully analyzed and showed comparable results to the established IPL method. The accuracy of the compartments identified with the proposed Python method was further corroborated based on the DSC values (Supplementary Figure 1A), which showed median values of 0.989, 0.991, and 1.000 for the trabecular, cortical, and full vertebra compartments, respectively. These values reflect minor differences, as shown in the relative difference between compartments (Supplementary Figure 1A). Crucially, the proposed implementation conserved the sensitivity to discriminate biological differences (Supplementary Figure 1B), as the IPL and Python implementations showed no significant differences over the four weeks while preserving the statistical difference observed between groups. Additionally, the Spearman correlation coefficients were very high for the static and dynamic parameters evaluated (Supplementary Figure 1C), ranging from 0.989 for bone volume fraction (BV/TV) to 1.000 for bone formation rate (BFR).

Concerning the execution time, the IPL method took an average of 10 minutes and 32 seconds (median: 8 min. and 56 sec., interquartile range: 8 min. 31 sec. – 10 min. 08 sec.) of CPU time per sample, running on HP OpenVMS (version V8.4) with HP Integrity BL890c i2 (1.60 GHz, 3 cores per running execution). Conversely, the Python implementation needed an average of 1 minute and 22 seconds (median: 1 min. and 22 sec., interquartile range: 1 min. 20 sec. – 1 min. 24 sec.) of CPU time per sample running on Fedora 35 with Intel Xeon Platinum 8168 CPU (2.7-3.7 GHz, up to 48 cores per running execution). Since the pipeline is faster, a higher number of samples can be processed in a very short period. Moreover, as each sample is analyzed in a single core, our parallel implementation further reduces the processing time, being only limited by the number of cores available. For instance, the images from a 4-week study comprising 40 samples per time-point would require four iterations on a 48-core CPU to perform this compartmental analysis, in less than 10 minutes, or the time equivalent for the IPL version to analyze a single sample. Even if the IPL implementation would process ten images in parallel, the proposed Python version would still represent at least a 16x speed up in processing time. More importantly, the proposed implementation can run on the desired HPC platform, ideally where subsequent micro-FE and post-processing steps will also be performed, reducing auxiliary data transfer operations to move data between systems, which would further increase the total overhead of the analysis and disk space requirements. Additionally, as the outputs generated for each sample are exported to a single file, the data management required to oversee all processed samples is considerably simplified compared to the IPL version.

##### 1.1.2 Micro-finite element analysis pipeline

As a second step, the micro-FE pipeline implemented in Python was also validated against the established method in IPL. Prior to the micro-FE, simulated intervertebral discs must be added at the top and bottom of the vertebra (Webster et al. 2008). The output of the proposed and established implementations achieved an average DSC of 0.997 (median: 0.998, interquartile range: 0.996 – 1.000). Furthermore, regarding the execution time for the default settings for all samples (stiffness: 14.8 GPa and radius: 13% of the height of the vertebra), the IPL implementation needed an average of 15.81 seconds (median: 16 sec., interquartile range: 15 sec. – 17 sec.) of CPU time per sample to add the intervertebral disc to the image, running on HP OpenVMS (version V8.4) with HP Integrity BL890c i2 (1.60 GHz). The proposed Python method required an average of 28.18 seconds (median: 27.60 sec., interquartile range: 26.64 sec. – 28.85 sec.) of CPU time per sample running on Fedora 35 with Intel Xeon Platinum 8168 CPU (2.7-3.7 GHz). Despite the slight increase in execution time, there are key advantages of the Python implementation, mainly that while the IPL version is designed to work exclusively on binary images, therefore only supporting its use for micro-FE analysis using homogeneous material properties, the proposed Python version expands this approach to grayscale images.

#### 1.2 Calibration of simulated intervertebral disc parameters

A preliminary analysis assessed the effect of the Young's modulus value and radius of the simulated intervertebral discs added to the vertebra on the ability to quantify mechanoregulation information from time-lapsed micro-CT data and identify the most suitable parameters to be used for the micro-FE analysis with homogeneous and heterogeneous material properties. The default radius on the established IPL pipeline is 13% of the height of the vertebra (Webster et al. 2008), and previous work has used stiffness values of 2.5 MPa (Webster et al. 2008) and 14.8 GPa (Schulte et al. 2013). Therefore, a disc radius between 7% and 17% of the height of the vertebra and ten stiffness values regularly spaced on a logarithmic scale between 2.5 MPa and 20 GPa (and including 14.8 GPa) were considered. Given the considerable amount of parameter combinations, only the samples of the static loading group from the first time-point were considered, as this is the *in vivo* group that best mimics the loading condition of the micro-FE analysis.

A micro-FE analysis was performed for all selected samples, using homogeneous and heterogeneous properties, and the mechanoregulation information was quantified using the correct classification rate (CCR) (Tourolle né Betts et al. 2020) as an indicator of the performance of each. The best-performing settings for each configuration were used for the mechanoregulation analysis performed in the remaining work.

For the micro-FE analysis with homogeneous material properties, the ability to retrieve mechanoregulation information from time-lapsed data showed slight variation (below 1%) across the domain space. Nonetheless, a stiffness value of 14.8 GPa (the same value assigned to bone) or higher showed slightly higher CCR values. Concerning the disc radius, a value between 11% and 13% of the vertebra height also matched the region of the highest CCR values. Additionally, this setting enabled the solver to converge for all samples in a single run, while lower disc stiffness values required adjusting the solver settings for some samples. Conversely, there was a higher variability of CCR values across the domain for heterogeneous material properties, with a stiffness of 18.4 MPa achieving the highest CCR value, for a radius of 13% of the vertebra height. In both instances, lower radii showed a decreasing trend in the amount of mechanoregulation information recovered. A critical remark is that the relationship applied to convert bone mineral density to Young's modulus yielded structures with an average stiffness ranging from 16.9 ± 4.3 GPa (week 0) to 17.5 ± 4.5 GPa (week 4), which is higher than the corresponding Young's Modulus value used for homogeneous material properties.

### 2 Supplementary Figures and Tables

#### 2.1 Supplementary Figures

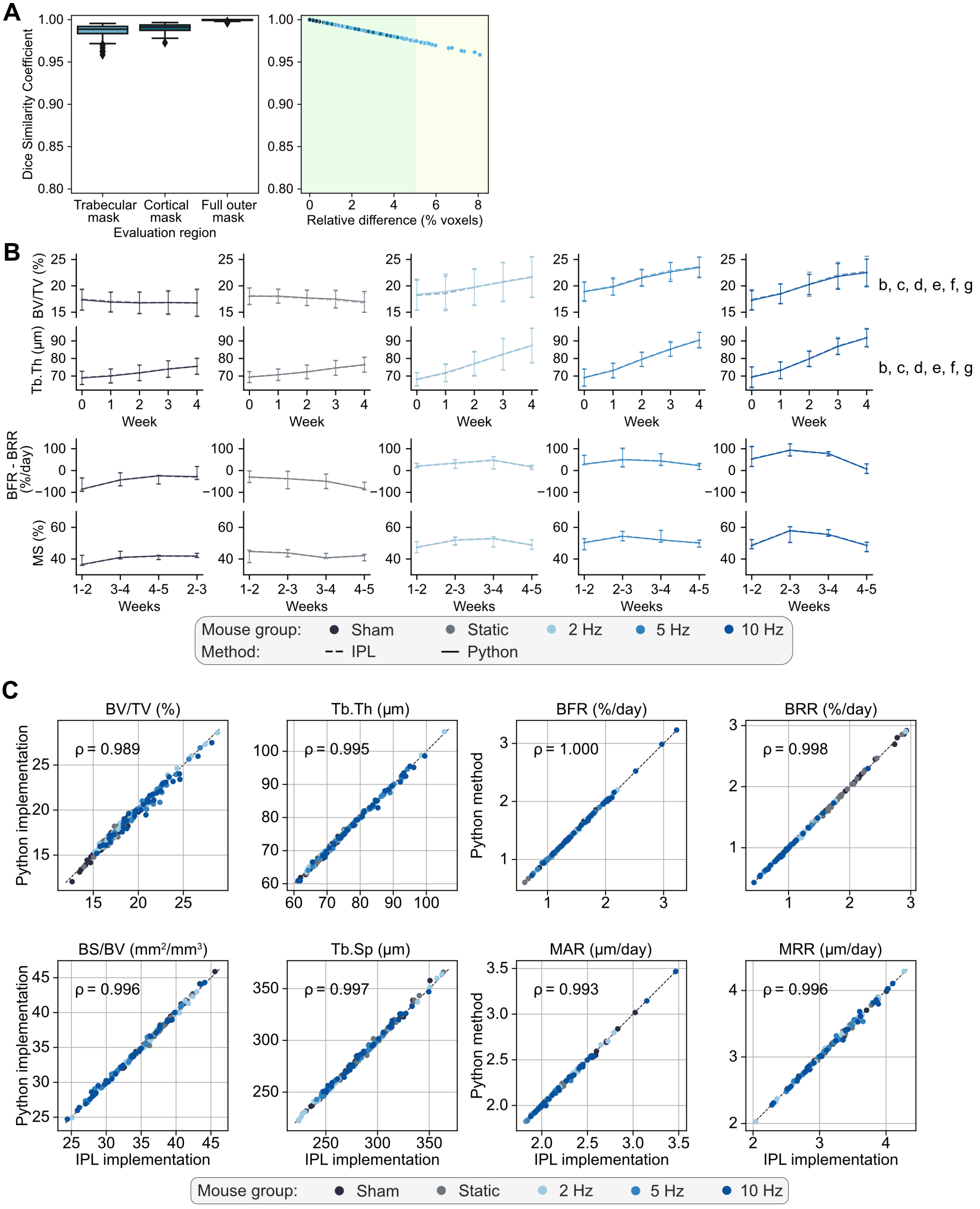

**Supplementary Figure 1.** Comparison of bone morphometric parameters computed for the regional compartments generated using the established IPL and the proposed Python implementations. A) Dice similarity coefficients computed between the compartments identified with each method for all samples and all time-points. Data is shown per compartment and based on the relative percentage difference of voxels between compartments, assuming the IPL method as the reference. B) Selected static and dynamic bone morphometric parameters computed for all time-points, using compartments identified with the IPL (dashed line) and proposed Python (full line) methods, showing no statistical differences between both methods ($p>0.05$), as determined by linear mixed effects model; furthermore, significant differences between groups determined by Tukey post-hoc multiple comparisons test of the values from the last timepoint ($p<0.05$, a: 2 Hz-5 Hz, b: 2 Hz-Sham, c: 2 Hz-Static, d: 5 Hz-Sham, e: 5 Hz-Static, f: 10 Hz-Sham, g: 10 Hz-Static) are still conserved for the proposed Python implementation. C) Correlation between static morphometry parameters (bone volume fraction - BV/TV, specific bone surface - BS/BV, trabecular thickness - Tb.Th, trabecular spacing - Tb.Sp) and dynamic morphometry parameters (bone formation rate - BFR, mineral apposition rate - MAR, bone resorption rate - BRR, mineral resorption rate - MRR) computed with each method for all samples and all time-points. The Spearman correlation coefficients computed show a near-perfect agreement in morphometric indices.

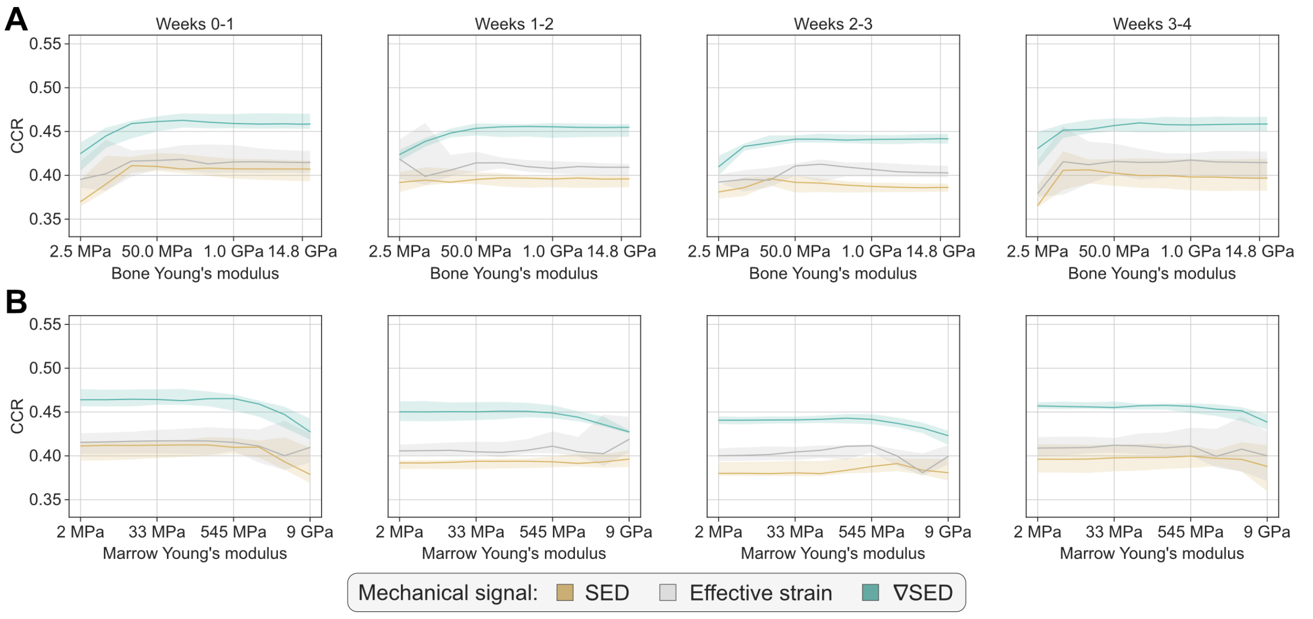


**Supplementary Figure 2.** Sensitivity analysis on the effect of the Young's modulus value used for bone and marrow voxels on the quantification of mechanoregulation information, using micro-FE analysis with homogeneous material properties. Only the static loading group was considered for this analysis. CCR values obtained by SED, effective strain, and ∇SED as local mechanical signal descriptors for the mechanoregulation analysis for weekly intervals. In A), bone tissue was assigned a Young's modulus varying between 2.5 MPa and 20.0 GPa, and marrow voxels (non-bone voxels in the trabecular region) were set to 2 MPa. Specifically, ten equally spaced values on a logarithmic scale between the two extremes were considered for the Young's modulus of bone (besides the original value of 14.8 GPa): 2.5 MPa, 6.8 MPa, 18.4 MPa, 50.0 MPa, 136.0 MPa, 368.0 MPa, 1.0 GPa, 2.7 GPa, 7.4 GPa, 14.8 GPa, 20.0 GPa. In B), bone tissue was assigned a Young's modulus of 14.8 GPa, and marrow voxels were assigned a value between 2.0 MPa (Webster et al. 2015) and 8.0 GPa, which reflects the range of Young's modulus observed in marrow voxels for the samples analyzed with heterogeneous material properties. Again, ten equally spaced values on a logarithmic scale between the two extremes were considered for the Young's modulus of marrow voxels: 2.0 MPa, 5.1 MPa, 13.0 MPa, 33.0 MPa, 84.1 MPa, 214.0 MPa, 545 MPa, 1.4 GPa, 3.5 GPa, 9.0 GPa. This analysis showed that the values selected in our study for the micro-FE analysis with homogeneous material properties (bone: 14.8 GPa, marrow: 2.0 MPa) do not influence the ability to retrieve mechanoregulation information from the data, given the uniform CCR values obtained.

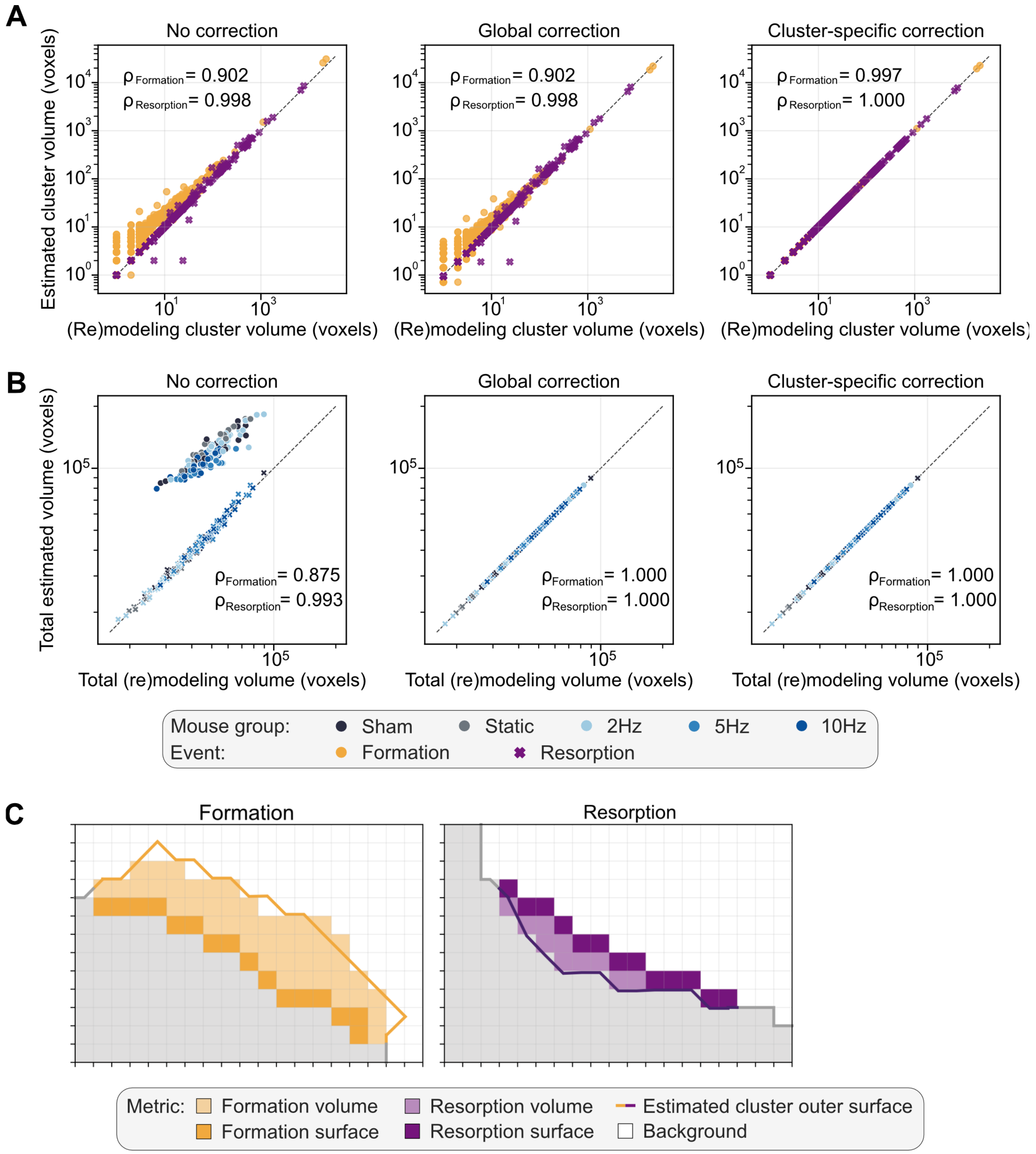


**Supplementary Figure 3.** A) Comparison between the estimated volume with the proposed mechanostat (re)modeling velocity estimation algorithm and actual volume measured by superimposing consecutive time-points for all (re)modeling clusters identified in one sample, with a 1-week interval. Three cases are shown: without any correction to the estimated (re)modeling distance (left), with a correction factor based on the total (re)modeling volume (middle), and with a correction factor specific to each (re)modeling cluster identified (right). The distance transform considered the taxicab metric. For each pair, the Spearman Correlation Coefficient is presented. B) Comparison of the total estimated volume with the proposed mechanostat (re)modeling velocity estimation algorithm and actual volume measured between time-points, for all samples and weekly intervals, including or not the correction factor depicted in A) for a single sample, with the corresponding Spearman correlation coefficient for each event. Without any correction, the (re)modeling distance collected from the (re)modeling surfaces yielded a volume change that did not match the volume of their corresponding clusters nor the total volume of (re)modeling clusters between time-points. Using a global correction, the total (re)modeled volume was accurately captured, but the estimated volume of individual (re)modeling clusters still did not match (A and B, middle), leading to a skewed pairing of remodeling distance and mechanical signal values. Finally, using a cluster-specific correction, the estimation obtained from surface voxels is scaled to reflect the corresponding cluster volume accurately for all samples. The volume of each cluster is computed, as well as the estimation from the surrounding surface voxels. Since the resolution of the micro-CT images does not allow determining with certainty from which voxel a given cluster was formed/resorbed, we assume that all neighboring surface voxels of the cluster contribute equally to that process, which is mathematically translated to a linear scaling of the (re)modeling distance values collected from the surface voxels associated with a given cluster to match the change required to produce the volume of the cluster. C) two-dimensional example of the outcome of the surface estimation performed with the proposed algorithm to compute (re)modeling distances for each surface voxel. Using a distance transform applied on the follow-up and inverse of the follow-up images, the displacement of the surface between time-points can be estimated for formation and resorption clusters to approximate the associated volume change of the (re)modeling cluster. Using the cluster-specific volume correction, we ensure that the predicted volume from the surface voxels matches the computed volume of the cluster. Both images were isolated from a three-dimensional micro-CT image and show how formation and resorption clusters look and how the corresponding surface is estimated with our approach..

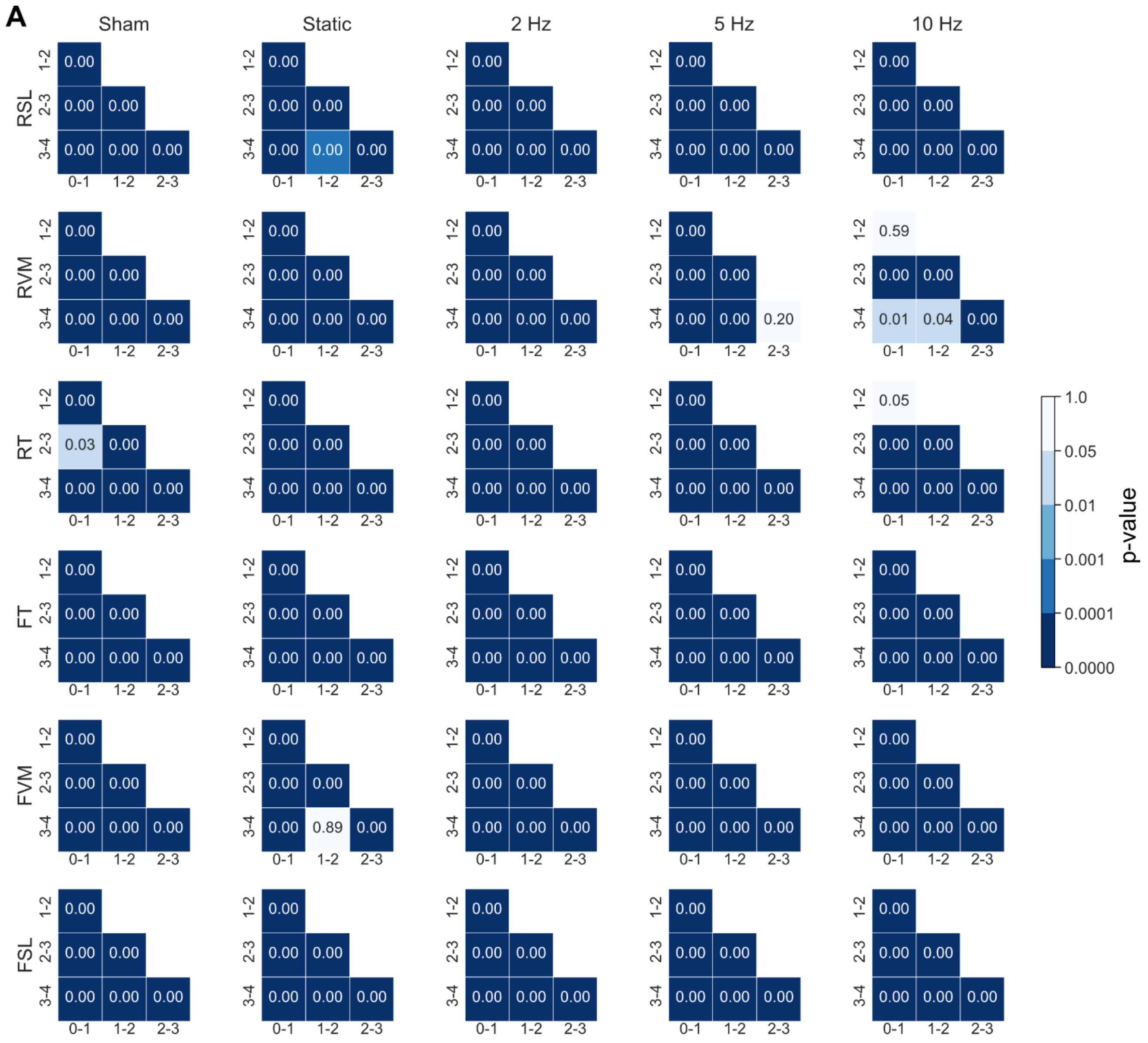


**Supplementary Figure 4.** A) Matrices with the p-values for multiple comparisons between week intervals (0-1, 1-2, 2-3, 3-4) of the parameter distributions obtained from the piecewise linear function for each group, estimated using the balanced bootstrapping approach described in the methods (see section “Mechanostat (re)modeling velocity curve and parameter derivation”). Parameter legend (see Materials and Methods and Table 1 for an extended description): Resorption saturation level (RSL), Resorption velocity modulus (RVM), Resorption threshold (RT), Formation threshold (FT), Formation velocity modulus (FVM), Formation saturation level (FSL). Statistical significance was determined with Conover’s test corrected for multiple comparisons with a step-down method using Bonferroni-Holm adjustments.

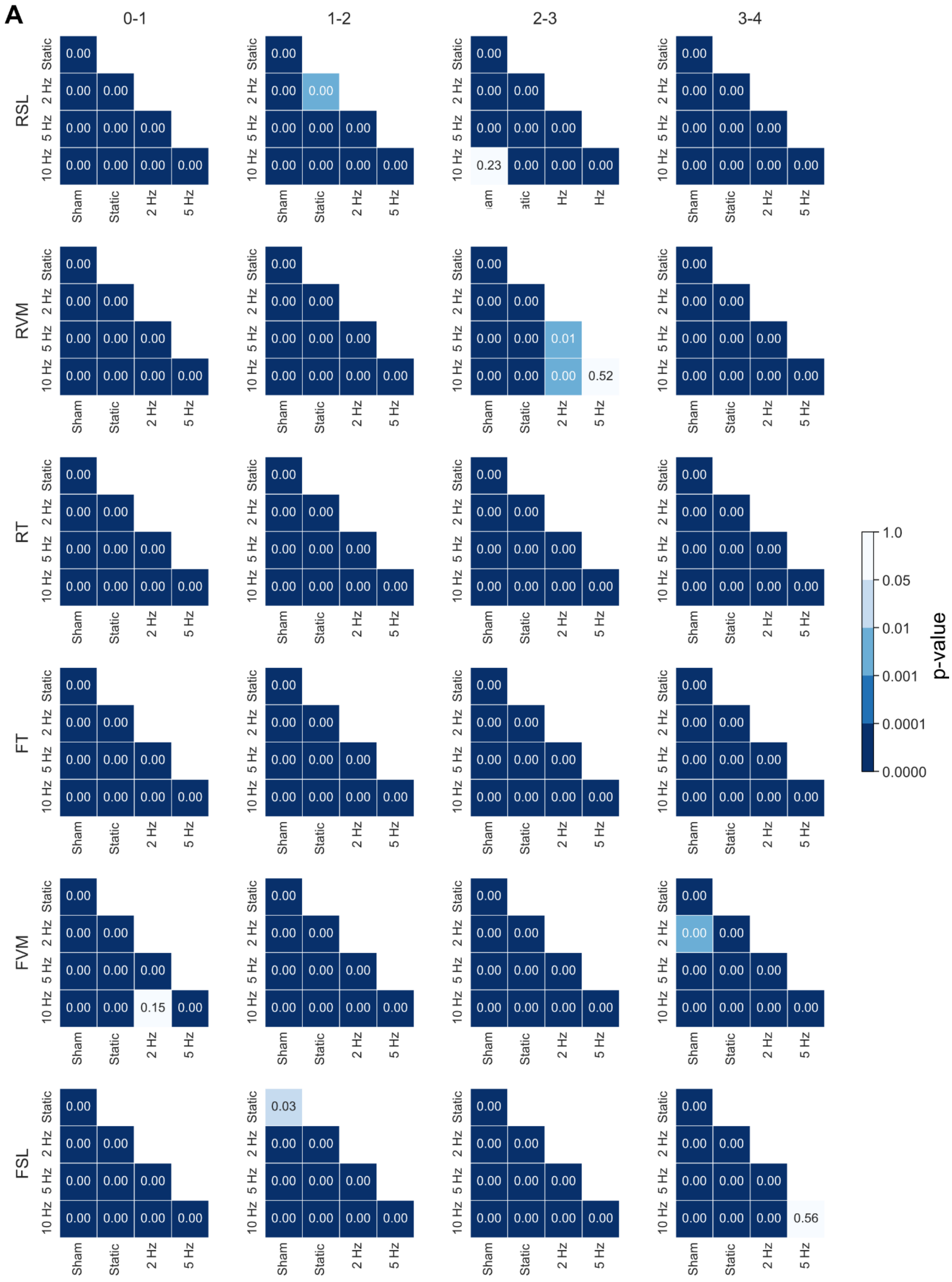
A
0-1
1-2
2-3
3-4
RSL
RVM
RT
FT
FVM
FSL
Static
2 Hz
5 Hz
10 Hz
Sham
p-value
1.0
0.05
0.01
0.001
0.0001
0.0000

**Supplementary Figure 5.** A) Matrices with the p-values for multiple comparisons between groups of the parameter distributions obtained from the piecewise linear function for each week interval (0-1, 1-2, 2-3, 3-4), estimated using the balanced bootstrapping approach described in the methods (see section "Mechanostat (re)modeling velocity curve and parameter derivation"). Parameter legend (see Materials and Methods and Table 1 for an extended description): Resorption saturation level (RSL), Resorption velocity modulus (RVM), Resorption threshold (RT), Formation threshold (FT), Formation velocity modulus (FVM), Formation saturation level (FSL). Statistical significance was determined with Conover's test corrected for multiple comparisons with a step-down method using Bonferroni-Holm adjustments.

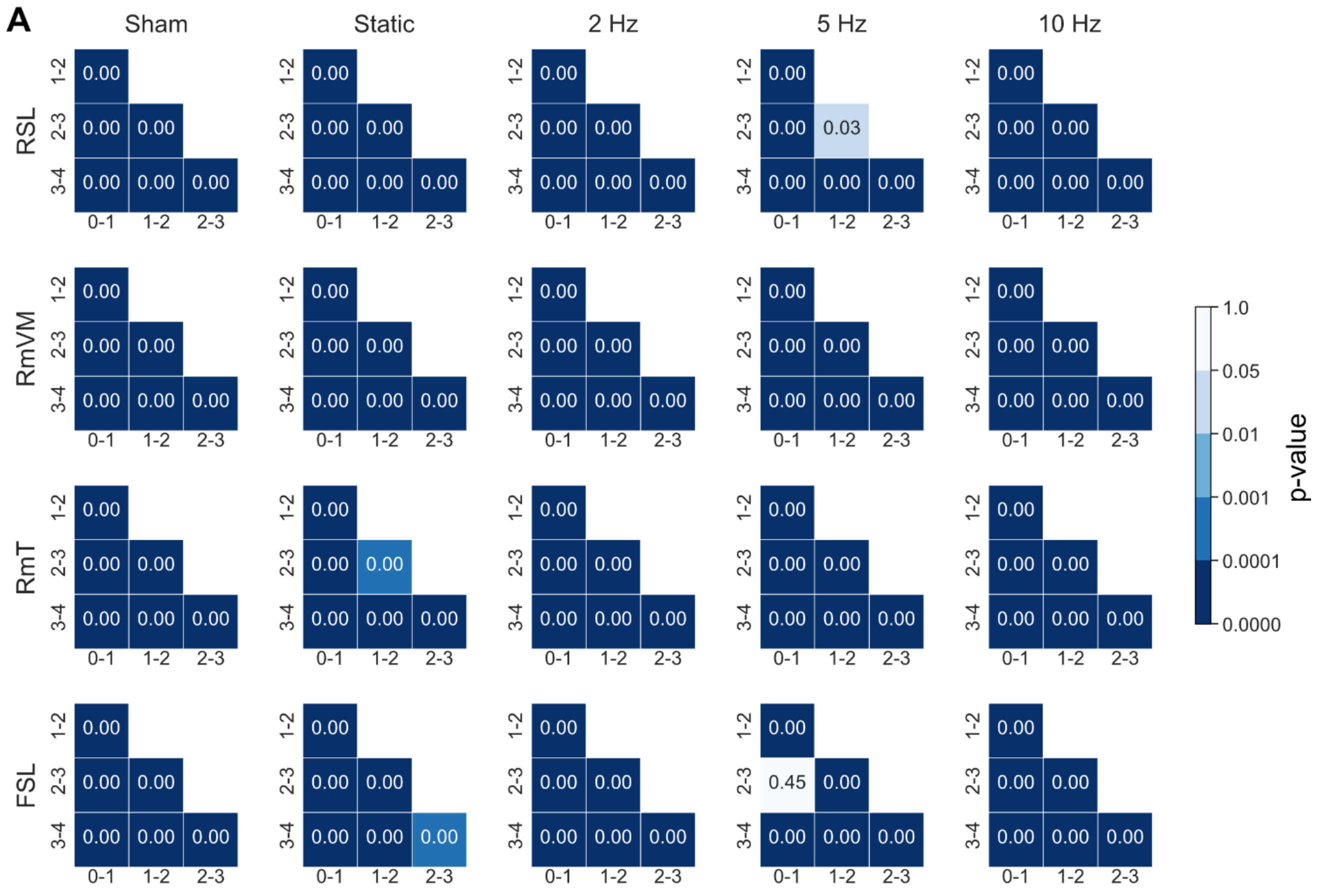


**Supplementary Figure 6.** A) Matrices with the p-values for multiple comparisons between week intervals (0-1, 1-2, 2-3, 3-4) of the parameter distributions obtained from the hyperbola function for each group, estimated using the balanced bootstrapping approach described in the methods (see section "Mechanostat (re)modeling velocity curve and parameter derivation"). Parameter legend (see Materials and Methods and Table 1 for an extended description): Resorption saturation level (RSL), (Re)modeling threshold (RmT), (Re)modeling velocity modulus (RmVM), Formation saturation level (FSL). Statistical significance was determined with Conover's test corrected for multiple comparisons with a step-down method using Bonferroni-Holm adjustments.

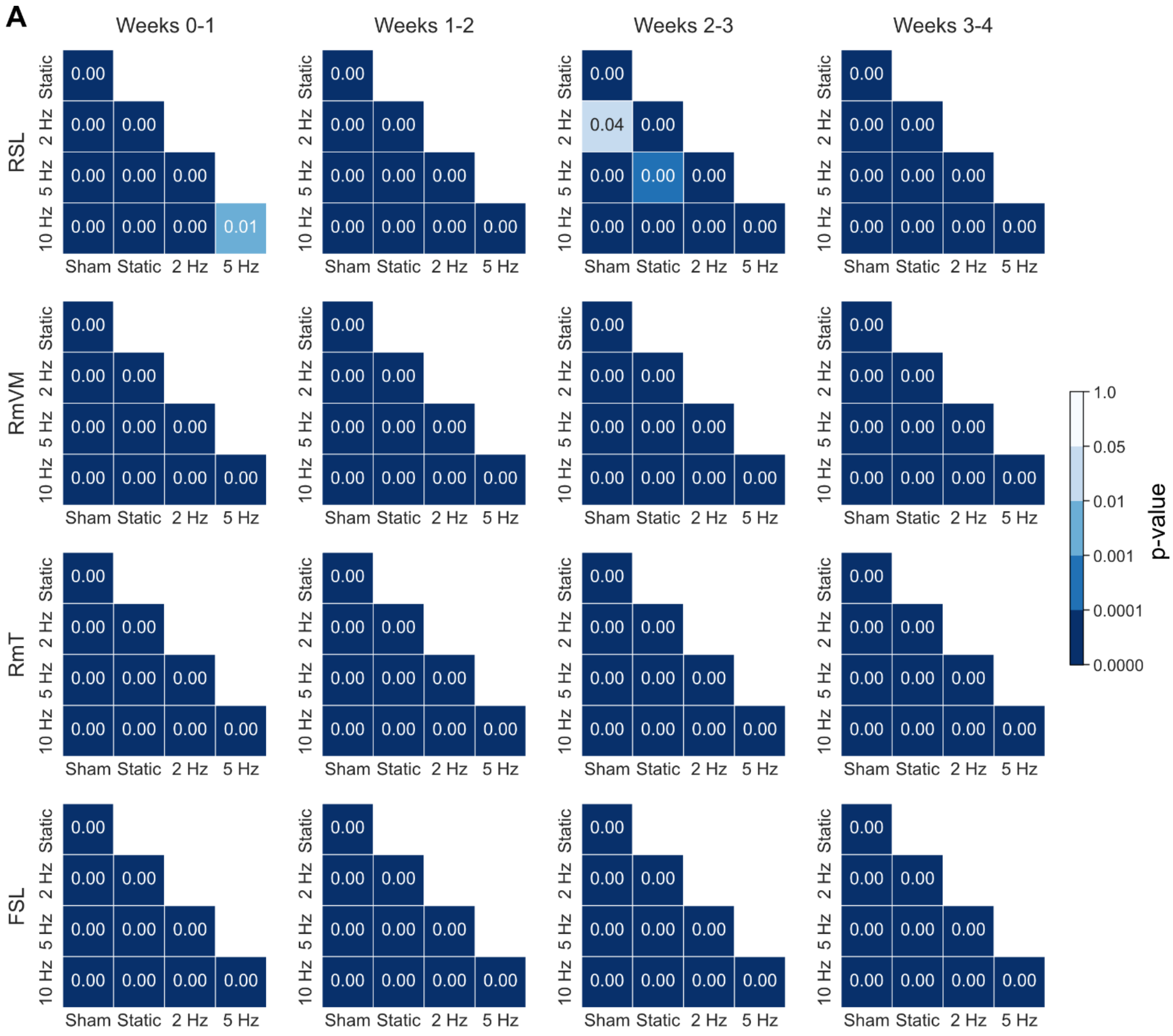


**Supplementary Figure 7.** A) Matrices with the p-values for multiple comparisons between groups of the parameter distributions obtained from the hyperbola function for each week interval (0-1, 1-2, 2-3, 3-4), estimated using the balanced bootstrapping approach described in the methods (see section "Mechanostat (re)modeling velocity curve and parameter derivation"). Parameter legend (see Materials and Methods and Table 1 for an extended description): Resorption saturation level (RSL), (Re)modeling threshold (RmT), (Re)modeling velocity modulus (RmVM), Formation saturation level (FSL). Statistical significance was determined with Conover's test corrected for multiple comparisons with a step-down method using Bonferroni-Holm adjustments.

#### 2.2 Supplementary Tables

**Supplementary Table 1:** Parameters of the mathematical functions fitted to the estimated mechanostat group average (re)modeling velocity curves for all weekly intervals. Data presented as "parameter (IQR)", where the IQR was calculated using the balanced bootstrapping approach described in the methods (see section "Mechanostat (re)modeling velocity curve and parameter derivation") to provide an estimate for the variability of each parameter. Root mean squared error (RMSE) was used to characterize the quality of the fit. The row "Effective strain range" indicates the range of mechanical signal values from which the fit of the mathematical functions was derived. Parameter legend (see Materials and Methods and Table 1 for an extended description): Resorption saturation level (RSL), Resorption velocity modulus (RVM), Resorption threshold (RT), Formation threshold (FT), Formation velocity modulus (FVM), Formation saturation level (FSL), (Re)modeling threshold (RmT), (Re)modeling velocity modulus (RmVM).

| Weeks | Function | Parameter | Unit | Group | | | | |
|---|---|---|---|---|---|---|---|---|
| | | | | Sham | Static | 2Hz | 5Hz | 10Hz |
| 0-1 | Piecewise linear | RSL | μm/day | -1.988 (-2.306 - -1.484) | -0.934 (-1.108 - -0.914) | -0.939 (-1.370 - -0.790) | -0.699 (-0.781 - -0.531) | -0.885 (-1.195 - -0.696) |
| | | RVM ($x10^3$) | (μm/day) / με | 12.832 (6.715 - 17.396) | 1.861 (1.708 - 2.837) | 2.683 (3.133 - 5.022) | 2.794 (1.744 - 4.964) | 6.398 (4.436 - 42.888) |
| | | RT | με | 155 (134 - 224) | 512 (397 - 556) | 360 (256 - 332) | 260 (193 - 342) | 148 (48 - 170) |
| | | FT | με | 490 (460 - 545) | 720 (630 - 750) | 370 (340 - 485) | 260 (260 - 390) | 180 (150 - 280) |
| | | FVM ($x10^3$) | (μm/day) / με | 0.306 (0.506 - 1.065) | 0.361 (0.313 - 0.364) | 0.487 (0.444 - 0.564) | 0.370 (0.425 - 0.554) | 0.468 (0.463 - 0.543) |
| | | FSL | μm/day | 0.220 (0.073 - 0.193) | 0.548 (0.280 - 0.460) | 0.515 (0.470 - 0.608) | 0.804 (0.433 - 0.561) | 1.146 (0.680 - 1.008) |
| | | RMSE | μm/day | 0.203 (0.191 - 0.247) | 0.126 (0.102 - 0.119) | 0.138 (0.099 - 0.148) | 0.111 (0.066 - 0.094) | 0.158 (0.123 - 0.165) |
| | Hyperbolic | RSL | μm/day | -2.956 (-3.312 - -2.574) | -1.048 (-1.404 - -1.140) | -1.082 (-1.592 - -0.754) | -0.583 (-0.765 - -0.554) | -0.392 (-0.916 - -0.297) |
| | | RmVM ($x10^{-3}$) | (μm/day) x με | -0.113 (-0.128 - -0.092) | -0.886 (-0.636 - -0.398) | -0.637 (-0.756 - -0.498) | -1.084 (-0.817 - -0.395) | -4.263 (-3.586 - -0.574) |
| | | RmT | με | 0.456 (0.410 - 0.623) | 0.700 (0.598 - 0.734) | 0.452 (0.391 - 0.478) | 0.408 (0.281 - 0.411) | 0.373 (0.246 - 0.378) |
| | | FSL | μm/day | 0.144 (0.059 - 0.156) | 0.424 (0.272 - 0.354) | 0.553 (0.493 - 0.590) | 0.663 (0.441 - 0.561) | 1.002 (0.637 - 0.914) |
| | | RMSE | μm/day | 0.097 (0.096 - 0.129) | 0.125 (0.082 - 0.100) | 0.133 (0.093 - 0.131) | 0.121 (0.067 - 0.092) | 0.175 (0.131 - 0.172) |
| | Effective strain range | | με | 1–1280 | 10–2240 | 10–2060 | 10–2520 | 10–2660 |
| 1-2 | Piecewise linear | RSL | μm/day | -2.234 (-2.951 - -1.623) | -1.506 (-1.676 - -1.460) | -1.6541 (-1.9388 - -1.3719) | -1.355 (-1.529 - -1.215) | -1.064 (-1.316 - -0.944) |
| | | RVM ($x10^3$) | (μm/day) / με | 34.990 (10.189 - 41.670) | 3.472 (3.244 - 4.503) | 8.567 (5.903 - 11.450) | 7.520 (7.641 - 10.713) | 8.104 (6.993 - 17.059) |
| | | RT | με | 74 (64 - 147) | 444 (360 - 483) | 203 (180 - 244) | 190 (163 - 209) | 141 (80 - 160) |

| | | | | | | | | |
|---|---|---|---|---|---|---|---|---|
| | | FT | με | 370 (360 - 410) | 746 (626 - 844) | 350 (320 - 390) | 220 (200 - 294) | 180 (110 - 190) |
| | | FVM (x10$^{3}$) | (μm/day) / με | 0.553 (0.504 - 0.663) | 0.245 (0.207 - 0.318) | 0.606 (0.549 - 0.662) | 0.685 (0.875 - 1.428) | 1.037 (0.904 - 1.090) |
| | | FSL | μm/day | 0.213 (0.196 - 0.276) | 0.327 (0.159 - 0.294) | 0.489 (0.458 - 0.542) | 0.588 (0.434 - 0.590) | 0.717 (0.657 - 0.727) |
| | | RMSE | μm/day | 0.222 (0.180 - 0.267) | 0.182 (0.168 - 0.214) | 0.124 (0.109 - 0.138) | 0.157 (0.094 - 0.113) | 0.144 (0.104 - 0.128) |
| | Hyperbolic | RSL | μm/day | -2.457 (-2.710 - -2.069) | -2.210 (-2.798 - -2.246) | -1.853 (-2.271 - -1.652) | -1.562 (-1.786 - -1.489) | -1.153 (-1.327 - -0.889) |
| | | RmVM (x10$^{-3}$) | (μm/day) x με | -0.120 (-0.145 - -0.104) | -0.332 (-0.330 - -0.216) | -0.318 (-0.374 - -0.259) | -0.284 (-0.280 - -0.226) | -0.296 (-0.360 - -0.246) |
| | | RmT | με | 338 (310 - 360) | 740 (641 - 901) | 361 (323 - 388) | 263 (221 - 314) | 194 (169 - 212) |
| | | FSL | μm/day | 0.213 (0.184 - 0.252) | 0.236 (0.139 - 0.236) | 0.517 (0.478 - 0.554) | 0.628 (0.507 - 0.679) | 0.774 (0.706 - 0.787) |
| | | RMSE | μm/day | 0.189 (0.126 - 0.230) | 0.140 (0.110 - 0.151) | 0.116 (0.093 - 0.122) | 0.146 (0.096 - 0.114) | 0.126 (0.091 - 0.113) |
| | Effective strain range | | με | 1–1150 | 10–2090 | 10–2000 | 10–2130 | 10–2150 |
| 2-3 | Piecewise linear | RSL | μm/day | -1.595 (-1.940 - -1.289) | -1.069 (-1.271 - -1.046) | -1.763 (-1.929 - -1.540) | -1.247 (-1.832 - -1.076) | -1.392 (-2.026 - -1.224) |
| | | RVM (x10$^{3}$) | (μm/day) / με | 9.198 (6.345 - 13.584) | 2.385 (1.986 - 3.393) | 13.608 (9.448 - 18.431) | 10.830 (7.819 - 20.311) | 10.027 (8.609 - 23.894) |
| | | RT | με | 173 (135 - 204) | 458 (379 - 547) | 140 (114 - 190) | 125 (89 - 143) | 149 (95 - 152) |
| | | FT | με | 350 (340 - 390) | 757 (683 - 867) | 221 (181 - 312) | 160 (140 - 210) | 200 (160 - 210) |
| | | FVM (x10$^{3}$) | (μm/day) / με | 0.619 (0.564 - 0.715) | 0.760 (0.200 - 0.484) | 0.637 (0.615 - 0.822) | 0.859 (0.826 - 1.186) | 1.152 (0.971 - 1.213) |
| | | FSL | μm/day | 0.216 (0.180 - 0.240) | 0.094 (0.075 - 0.196) | 0.411 (0.357 - 0.459) | 0.476 (0.424 - 0.508) | 0.545 (0.537 - 0.586) |
| | | RMSE | μm/day | 0.139 (0.117 - 0.177) | 0.156 (0.122 - 0.144) | 0.141 (0.116 - 0.152) | 0.120 (0.081 - 0.103) | 0.144 (0.118 - 0.173) |
| | Hyperbolic | RSL | μm/day | -2.223 (-2.688 - -1.709) | -1.527 (-1.890 - -1.545) | -2.105 (-2.356 - -1.834) | -1.464 (-2.053 - -1.248) | -1.701 (-2.549 - -1.385) |
| | | RmVM (x10$^{-3}$) | (μm/day) x με | -0.134 (-0.167 - -0.115) | -0.212 (-0.285 - -0.210) | -0.158 (-0.230 - -0.136) | -0.146 (-0.170 - -0.116) | -0.1938 (-0.239 - -0.153) |
| | | RmT | με | 0.337 (0.321 - 0.365) | 0.831 (0.639 - 0.955) | 0.254 (0.214 - 0.323) | 0.188 (0.161 - 0.225) | 0.207 (0.177 - 0.230) |
| | | FSL | μm/day | 0.234 (0.191 - 0.258) | 0.134 (0.112 - 0.223) | 0.442 (0.395 - 0.488) | 0.514 (0.465 - 0.550) | 0.600 (0.580 - 0.637) |
| | | RMSE | μm/day | 0.076 (0.066 - 0.092) | 0.149 (0.089 - 0.116) | 0.128 (0.098 - 0.138) | 0.113 (0.064 - 0.090) | 0.135 (0.102 - 0.149) |
| | Effective strain range | | με | 1–1170 | 10–2200 | 10–2030 | 10–1920 | 10–1940 |
| 3-4 | Piecewise linear | RSL | μm/day | -2.446 (-2.647 - -2.158) | -1.403 (-1.747 - -1.387) | -0.892 (-1.104 - -0.726) | -1.756 (-2.450 - -1.497) | -2.138 (-2.510 - -1.815) |
| | | RVM (x10$^{3}$) | (μm/day) / με | 28.921 (22.144 - 34.492) | 2.409 (2.356 - 4.024) | 3.430 (3.406 - 5.749) | 12.209 (9.288 - 20.387) | 11.534 (8.065 - 17.119) |
| | | RT | με | 85 (75 - 104) | 593 (437 - 607) | 270 (196 - 270) | 154 (114 - 195) | 195 (153 - 250) |
| | | FT | με | 290 (250 - 310) | 840 (770 - 1056) | 311 (240 - 360) | 340 (284 - 381) | 420 (340 - 420) |

| | | | | | | | | |
|---|---|---|---|---|---|---|---|---|
| | | FVM (x10³) | (μm/day) / με | 0.915 (0.681 - 0.963) | 0.298 (0.177 - 0.338) | 0.816 (0.732 - 0.866) | 0.825 (0.730 - 0.948) | 0.863 (0.636 - 0.813) |
| | | FSL | μm/day | 0.209 (0.223 - 0.300) | 0.096 (0.119 - 0.226) | 0.367 (0.332 - 0.445) | 0.411 (0.417 - 0.465) | 0.402 (0.397 - 0.498) |
| | | RMSE | μm/day | 0.176 (0.148 - 0.181) | 0.178 (0.154 - 0.185) | 0.132 (0.102 - 0.138) | 0.161 (0.122 - 0.186) | 0.184 (0.156 - 0.205) |
| | Hyperbolic | RSL | μm/day | -3.177 (-3.368 - -2.864) | -2.213 (-2.463 - -2.110) | -1.108 (-1.361 - -0.908) | -2.312 (-2.952 - -1.895) | -2.666 (-3.087 - -2.202) |
| | | RmVM ($x10^{-3}$) | (μm/day) x με | -0.083 (-0.101 - -0.084) | -0.337 (-0.407 - -0.321) | -0.245 (-0.341 - -0.243) | -0.168 (-0.215 - -0.162) | -0.256 (-0.368 - -0.215) |
| | | RmT | με | 254 (206 - 279) | 1030 (822 - 1092) | 314 (269 - 362) | 276 (246 – 317) | 372 (323 - 428) |
| | | FSL | μm/day | 0.231 (0.224 - 0.297) | 0.153 (0.133 - 0.222) | 0.427 (0.403 - 0.504) | 0.427 (0.425 - 0.459) | 0.446 (0.428 - 0.513) |
| | | RMSE | μm/day | 0.122 (0.074 - 0.092) | 0.148 (0.102 - 0.131) | 0.142 (0.105 - 0.138) | 0.131 (0.087 - 0.126) | 0.158 (0.125 - 0.169) |
| | Effective strain range | | με | 1–1200 | 10–2310 | 10–2000 | 10–1810 | 10–1830 |

**Supplementary Table 2:** Goodness of fit of a logarithmic function to the mechanostat parameters estimated from RmV curves for each loading frequency. Parameter distributions are bootstrapped per group, and the median value was selected. The parameters “y0” and “a” define the logarithmic function considered (see Materials and Methods), with the units of each parameter. Pseudo-$R^2$ is used to quantify the quality of the fit. Parameter legend (see Materials and Methods and Table 1 for an extended description): Resorption saturation level (RSL), Resorption velocity modulus (RVM), Resorption threshold (RT), Formation threshold (FT), Formation velocity modulus (FVM), Formation saturation level (FSL), (Re)modeling threshold (RmT), (Re)modeling velocity modulus (RmVM).

| Piecewise linear function | | | | |
|---|---|---|---|---|
| Weeks | Parameter name | y0 | a | Pseudo-$R^2$ |
| 0-1 | RSL | -0.87855 | 0.05945 | 0.297 |
| | RVM | Did not converge | | |
| | RT | 0.00033 | -0.00008 | **0.966** |
| | FT | 0.00047 | -0.00011 | **0.994** |
| | FVM | 416.18299 | 36.93416 | **0.897** |
| | FSL | 0.48264 | 0.05150 | 0.536 |
| 1-2 | RSL | -1.43046 | 0.07868 | 0.625 |
| | RVM | Did not converge | | |
| | RT | 0.00027 | -0.00007 | **0.989** |
| | FT | 0.00045 | -0.00013 | **0.997** |
| | FVM | Did not converge | | |
| | FSL | 0.43110 | 0.10244 | **0.962** |
| 2-3 | RSL | -1.44534 | -0.13352 | **0.714** |
| | RVM | 8986.16320 | 2973.86477 | **0.965** |
| | RT | 0.00023 | -0.00006 | **0.917** |
| | FT | 0.00031 | -0.00006 | 0.686 |
| | FVM | Did not converge | | |
| | FSL | 0.33400 | 0.09635 | **0.996** |
| 3-4 | RSL | -1.42008 | -0.00014 | 0.000 |
| | RVM | 7348.26286 | 1977.33847 | **0.700** |
| | RT | 0.00030 | -0.00007 | **0.942** |
| | FT | 0.00036 | -0.00001 | 0.058 |
| | FVM | Did not converge | | |
| | FSL | 0.32888 | 0.06941 | **0.984** |
| 0-4 | RSL | -2.74338 | 0.19865 | -0.060 |
| | RVM | Did not converge | | |
| | RT | 0.00031 | -0.00009 | **0.995** |
| | FT | 0.00039 | -0.00012 | 0.560 |
| | FVM | Did not converge | | |

| | | | | |
|---|---|---|---|---|
| | FSL | 1.15286 | 0.27029 | **0.987** |

| Hyperbola function | | | | |
|---|---|---|---|---|
| Weeks | Parameter name | y0 | a | Pseudo-$R^2$ |
| 0-1 | RSL | -0.91525 | 0.16595 | **0.800** |
| | RmVM | -0.00056 | -0.00003 | 0.148 |
| | RmT | 0.00049 | -0.00008 | **0.999** |
| | FSL | 0.46805 | 0.06973 | **0.820** |
| 1-2 | RSL | -1.99128 | 0.29318 | **0.956** |
| | RmVM | -0.00027 | 0.00000 | 0.061 |
| | RmT | 0.00044 | -0.00011 | **0.981** |
| | FSL | 0.43888 | 0.12305 | **0.989** |
| 2-3 | RSL | -1.81458 | -0.03794 | 0.120 |
| | RmVM | -0.00019 | 0.00002 | **0.727** |
| | RmT | 0.00022 | -0.00001 | 0.119 |
| | FSL | 0.37164 | 0.09942 | **0.994** |
| 3-4 | RSL | -1.86482 | 0.14229 | -0.086 |
| | RmVM | -0.00027 | 0.00004 | **0.657** |
| | RmT | 0.00028 | 0.00002 | -0.177 |
| | FSL | 0.34081 | 0.06702 | **0.958** |
| 0-4 | RSL | -2.99399 | 0.87960 | **0.756** |
| | RmVM | -0.00094 | -0.00008 | 0.000 |
| | RmT | 0.00044 | -0.00024 | 0.000 |
| | FSL | 1.16409 | 0.28317 | **0.911** |
